# Structural and functional characterisation of the dextran utilisome from *Bacteroides thetaiotaomicron*

**DOI:** 10.64898/2026.02.19.706672

**Authors:** Matthew Feasey, Augustinas Silale, Arnaud Basle, Bert van den Berg

## Abstract

*Bacteroides thetaiotaomicron* (*B. theta*) is a model Bacteroidota of the healthy human gut microbiota and a specialist in glycan utilisation. Like other *Bacteroides*, *B. theta* has many highly regulated polysaccharide utilisation loci (PUL) that encode outer membrane (OM) TonB-dependent transporters (SusC), closely associated “lid” lipoproteins (SusD), and additional surface-exposed lipoproteins (SLPs) that bind and partially degrade specific glycans derived from host cells, diet, or other microbiota members. The canonical starch PUL products are thought to form a dynamic complex in the presence of starch. However, other PULs form stable complexes in the absence of substrate (recently named “utilisomes”), with additional surface lipoproteins tightly associated with the core SusCD complex. In this study, we characterised the *B. theta* dextran utilisome, with a SusCD^dex^ core and an associated glycoside hydrolase (GH^dex^) and surface glycan binding protein (SBGP^dex^). Via X-ray crystallography we solved high-resolution structures of SBGP^dex^ in isolation and SusD^dex^ and GH^dex^ bound to dextran oligosaccharides. We used isothermal titration calorimetry (ITC) to quantify ligand binding of wild type and mutant SLPs. We further used single particle cryo-EM of the catalytically inactive dextran utilisome to visualise open and closed states of the complex. Three occupied dextran binding sites were observed across SusC^dex^, SusD^dex^ and GH^dex^, with substrate observed in both open and closed states of SusD^dex^. 3D variability analysis showed a minority of particles in the process of SusD^dex^ lid closure. Together our work defines commonalities and differences across utilisomes dedicated to the import of simple glycans.

## Introduction

The human gastrointestinal (GI) tract is home to a wide array of microbes, collectively termed the human gut microbiota (HGM). Intense research efforts over the past 2 decades have revealed that this bacteria-dominated community has far-reaching effects on human health. Complex dietary glycans (“fibre”) that cannot be degraded by enzymes of the human digestive tract are thought to be the primary nutrient source for the HGM, and the utilization of these complex sugars is crucial for the relationship between host and bacteria, leading, for example, to the generation of short-chain fatty acids that provide metabolic energy and systemic health benefits to the host. The colon or large intestine is the most densely populated part of the GI tract, with microbial densities up to 10^11^-10^12^ per gram of luminal content [1]. Most of the colonic microbiota belongs to two bacterial phyla: the Gram-positive Bacillota and the Gram-negative Bacteroidota. Additional, minor phyla include Actinomycetota and Pseudomonadota [1–3].

Belonging to the Bacteroidota, members of the genus *Bacteroides* have attracted particular attention for several reasons. First, several *Bacteroides* species (*B. fragilis*, *B. thetaiotaomicron*) [4,5] are relatively easy to culture in the lab and can be manipulated genetically. Second, the *Bacteroides* are collectively the most abundant bacterial genus in the gut of western world populations, comprising up to 40% of all bacteria in the large intestine [6]. As such, *Bacteroides* play an important role in shaping the HGM, and their study can be expected to yield important discoveries on fundamental aspects of the gut microbial ecosystem. Third, Bacteroides are prolific, primary degraders of many dietary glycans, producing many oligosaccharides that can be used by other members of the HGM, a phenomenon called cross-feeding. The best-studied member of the *Bacteroides*, *B. thetaiotaomicron* VPI-5482 *(B. theta)*, is considered a glycan “generalist” owing to the ∼18% of its 6.2 Mb genome that is dedicated to the utilisation of glycans [3,7].

The co-transcribed genes involved in the degradation of a particular glycan are organised on the genome in so-called PULs (polysaccharide utilisation loci). According to the PUL database (http://www.cazy.org/PULDB/), *B. theta* has 86 predicted PULs, and for most of these no ligands have been identified experimentally. PUL prediction is based on the presence of at least one SusCD pair, consisting of a SusC (from <u>s</u>tarch <u>u</u>tilisation <u>s</u>ystem or <u>s</u>accharide <u>u</u>tilisation <u>s</u>ystem) TonB-dependent transporter (TBDT) and a tightly associated SusD ligand binding protein. SusDs are surface-exposed lipoproteins (SLPs) that form a mobile cap on the SusC transporter to deliver transport-competent substrates to the SusC via a “pedal-bin” mechanism [8–10]. In addition to SusD proteins, PULs typically include additional SLPs such as surface glycan binding proteins (SGBPs) and glycoside hydrolases (GHs), involved in the initial binding of polymeric glycans and degradation of those glycans into transport-competent oligosaccharides, respectively [11,12]. It should be noted that approximately a quarter of predicted PULs do not contain proteins with obvious glycan binding or degradation functionalities, suggesting that non-glycans such as peptides [8] are also taken up by “PULs”.

Recently it was shown that for the “simple” glycan levan (a β2,6-linked fructan), all four outer membrane (OM) components (Bt1760-3) of the respective PUL form large (∼600 kDa) and stable complexes on the cell surface, irrespective of bound glycans, that were named “utilisomes” [10]. Cryo-EM structures of the purified utilisome revealed a dimer of tetramers each consisting of a GH^lev^, SGBP^lev^, SusD^lev^ and SusC^lev^, and the effect of levan substrates on the dynamics of the catalytically-inactivated utilisome was determined. In the same study, a cryo-EM structure of the wild-type, apo dextran utilisome was also reported [10] that showed a similar architecture as the levan utilisome, but with no information on substrate binding by any of the components. The dextran PUL (PUL48) is known to be upregulated in the presence of dextran [10,13,14].

Here, PUL48 was further characterised. We determined X-ray crystal structures of *E. coli*-expressed soluble variants of GH^dex^, SGBP^dex^, and SusD^dex^ without (SGBP^lev^) and with (GH^dex^ and SusD^dex^) bound dextran oligosaccharides, allowing the identification of important residues in GH^dex^ and SusD^dex^ for dextran binding. The affinities of the three proteins (wild type and binding mutants) for differently sized (∼1.5-500 kDa) dextran oligos were measured using isothermal titration calorimetry (ITC). For both GH^dex^ and SusD^dex^, the affinities for dextran are in the low μM range and roughly independent of dextran length. By contrast, the affinity of SGBP^dex^ for dextran oligos increases greatly with increasing ligand size to reach a similar affinity as GH^dex^ and SusD^dex^ for the longest dextran tested (500 kDa). The entire catalytically inactivated dextran utilisome was purified from *B. theta* cells via His-tagged SusD^dex^. Cryo-EM structures were determined in the presence of dextran substrate, revealing both similarities and differences compared to the levan utilisome that increase our understanding of glycan uptake by gut *Bacteroides*.

## Results

### Expression and purification of the dextran SLPs GH^dex^, SGBP^dex^ and SusD^dex^ from *E. coli*

For further analysis of the OM components of PUL48, GH^dex^, SGBP^dex^ and SusD^dex^ (*bt3087*, *bt3088* and *bt3089* respectively; Figure 1A), we generated *E. coli* constructs in which the cysteine that would be lipidated in the mature lipoprotein was removed. An additional 9 N-terminal residues, predicted by AlphaFold2 [15] to be unstructured, were also removed (Methods). This allowed high-yield overexpression of the PUL48 SLPs as soluble proteins. These proteins were then purified (Figure 1B) for X-ray crystallography and ITC. Average protein yields of the GH^dex^ constructs, including the catalytic mutants, was approximately 3 mg protein per litre of culture. Yields for SGBP^dex^ and SusD^dex^ were 7 and 5 mg/L, respectively.

**Figure 1.**
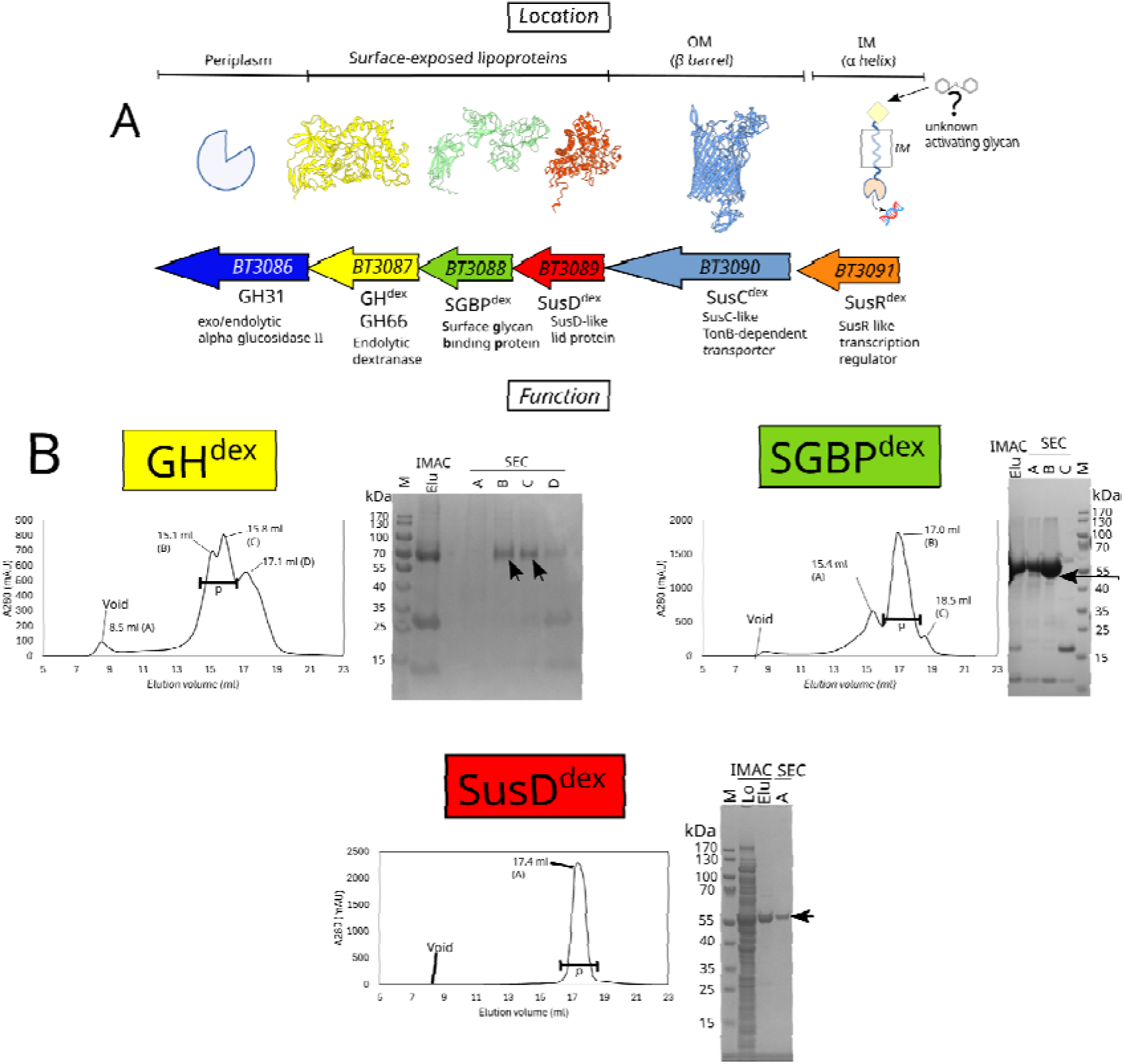
The dextran PUL of *B. theta* (PUL48) and recombinant expression of its SLPs. **A.** The dextran PUL and its constituent proteins, including Alphafold2 [15] models of the outer membrane associated components. **B**. Superose 6 SEC (size exclusion chromatography, Superose 6) chromatograms of GH^dex^, SGBP^dex^, and SusD^dex^ after recombinant expression in *E. coli* and IMAC (immobilised metal affinity chromatography, nickel affinity) purification. Pooled fractions are marked with “p” on the chromatograms. For GH^dex^, peaks B and C were pooled, while for SGBP^dex^ only peak B was pooled. SDS-PAGE (sodium dodecyl sulphate–polyacrylamide gel electrophoresis) gels of IMAC samples (either loaded sample (Lo) or final elutions (Elu)) and selected SEC fractions of each peak were loaded. Arrows indicate the final protein in the gel images. SEC samples were labelled corresponding to peak letters. Note that for SEC, Superose 6 columns had to be used since all the dextran binding proteins bind to (dextran-based) Superdex SEC resins.

### Structure determination of GH^dex^ in the absence and presence of substrate

GH^dex^ is a GH66 family endo-dextranase that hydrolyses the α1,6 backbone of dextran [16]. A study previously identified its two putative conserved catalytic residues, Asp297 and Glu360, by homology analysis between GH66 dextranases and chemical rescue experiments of *Paenibacillus* dextranase [16,17]. Homologous residues are also found in dextranases from *Streptococcus mutans* INGBRITT and *Bacillus circulans* T-3040 [16]. In the retaining mechanism of hydrolysis, the Asp carboxylate acts as a nucleophile while the Glu acts as an acid/base [16,18]. To verify the effect of the proposed GH^dex^ catalytic pair, we generated GH^dex^ pET28b constructs for D297A and E360A, as well as a double mutant D297A, E360A (GH^dex*^). These were then expressed as soluble proteins in *E. coli*. Since we assumed that GH^dex*^ would be completely inactive, we tested the E360A mutant for its ability to digest dextran 500 *in vitro* (Figure S1). WT GH^dex^ generated a ladder of glycans over the course of the reaction, culminating in glycans with a degree of polymerisation (DP) of 1-3. GH^dex^ E360A generated no degradation products, but the intensity of the time zero glycan spot decreased over time, suggesting some residual activity.

The X-ray structure of wild-type GH^dex^ was solved to a resolution of 1.8 Å (PDB: 9SJG), with apo GH^dex^ E360A solved to 2.1 Å (9SJH) and the same mutant co-crystallised with dextran 1.5 to 2.2 Å (9SJF) (see Figure 2, and Table S1 for X-ray crystallography statistics). GH^dex^ contains three domains. The N-terminal and C - terminal domains (NTD, CTD) contain Ig-like folds with 8 and 11 β-strands, respectively. Preceding the NTD, a short β-strand (residues 31-33) forms an inter-domain β-sheet with the CTD, which may rigidify these domains. The catalytic domain consists of a (β/α)_8_ barrel (residues 136-469, Figure 2B), and the catalytic pair D297 and E360 [19,20] is found within the catalytic domain cavity on the 4^th^ and 6^th^ β-strands of the barrel. The catalytic domain forms a pocket or crater-like topology, which has been previously described in glycoside hydrolases that hydrolyse exopolysaccharides [21]. The distance between the catalytic carboxylates is 6.5 Å, similar to the reported distance in retaining mechanism enzymes [18,21].

**Figure 2.**
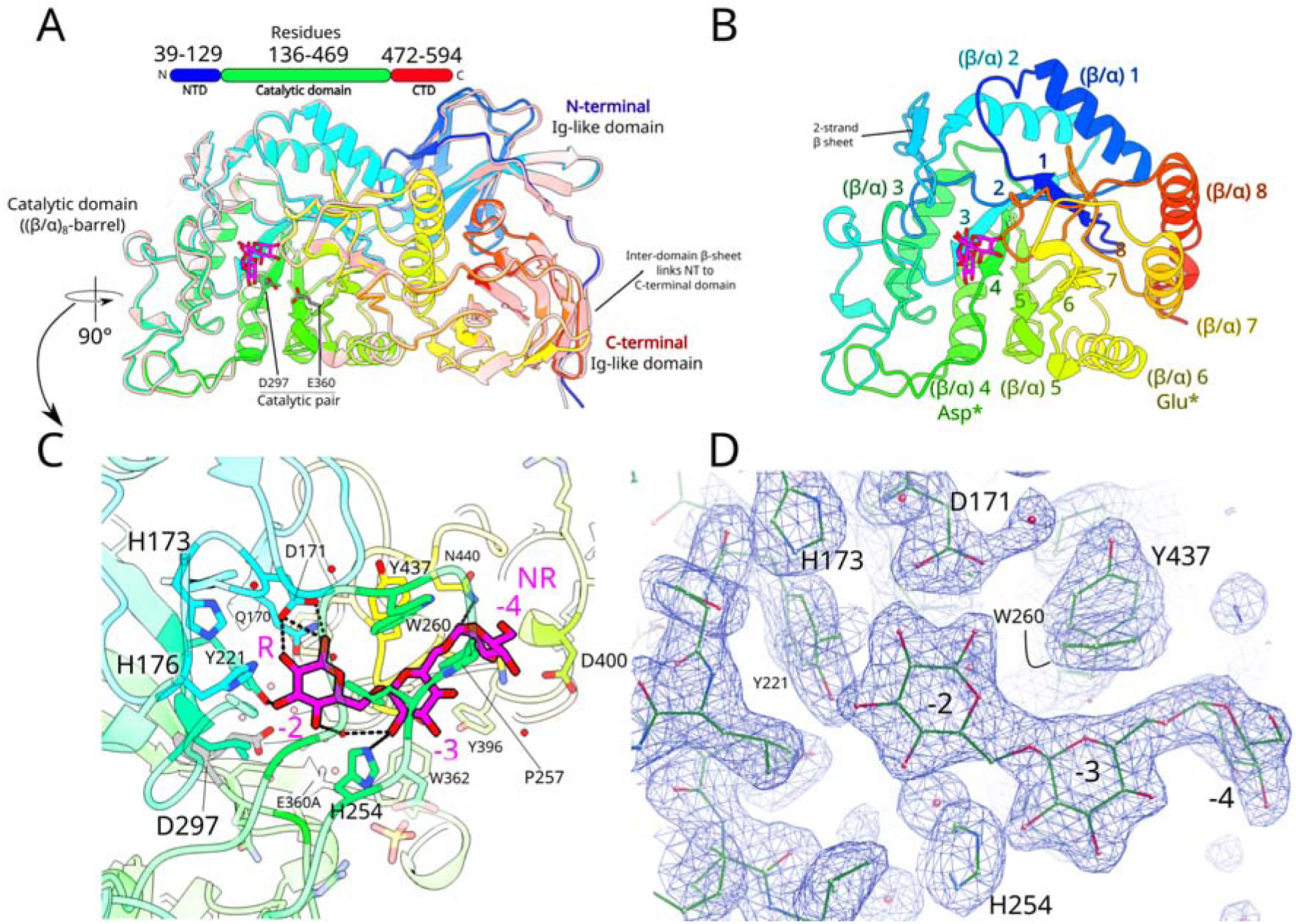
The crystal structures of wild-type (WT) GH^dex^ and the catalytic mutant GH^dex^ E360A. **A**. Overlaid cartoon displays of apo WT GH^dex^ (in salmon) and the E360A catalytic inactive mutant (in rainbow) with bound dextran. Domain assignment is shown for the primary sequence at top, coloured by domain (N-terminal domain (NTD) is in blue, the catalytic domain is in green, and the C-terminal domain (CTD) is in red). In the WT structure the side chains of the catalytic residues D297 and E360 are displayed in grey. **B**. The (β-α)_8_ catalytic domain of GH^dex^ E360A is shown and recoloured in rainbow format. β-strands that contain the catalytic residues are marked with an asterisk. **C**. The binding site of GH^dex^. Putative subsites for each glucose residue are numbered, with non-reducing (NR) and reducing (R) ends labelled. Residues were first numbered (with increasingly negative numbers from the catalytic site to the NR end) based on homology with TpDex [20] (Figure S2) and then confirmed in the cryo-EM structure of GH^dex^ (Figure 5). Residues < 5 Å from the dextran ligand are highlighted in full colour, with side chains displayed for residues with hydrogen bonds to the ligand (represented by dashed lines), residues that could hydrogen bond with a longer ligand, and aromatic residues likely important for binding. The catalytic residues (D297, E360A) are coloured in grey. **D.** The modelled dextran is shown in the 2Fo-Fc map at 1 σ in Coot. Structures of apo GH^dex^ (WT) and GH^dex^ E360A with and without dextran are available with accession codes 9SJG, 9PJF and 9SJH, respectively.

As expected, GH^dex^ is divergent from GH^lev^ (GH32) and the canonical SusG (GH^starch^, GH13) [10,22], with Cα RMSD values of 14.3 Å and 48.5 Å, respectively. Comparison with the GH^dex^ Alphafold prediction indicated minimal differences with a Cα RMSD of 0.6 Å. When GH^dex^ was compared to *Thermoanaerobacter pseudoethanolicus* dextranase (*Tp*^dex^, 5AXG, 5AXH [20], see also Figure S2 and the Supplementary discussion), the structures were found to be highly similar with Cα RMSD of 3.2 Å, or 1.0 Å when only the catalytic domains were compared. In the mutant *Tp*^dex^ structure liganded with isomaltohexaose, the authors replaced the catalytic nucleophile (D312G), and the bound ligand spanned subsites −4 to +2. Comparison with GH^dex^ E360A showed that the ligand conformations were highly similar, with three glucose residues of dextran 1.5 resolved above the active site (subsites −2 to −4). This suggests that the E360A mutant retained some catalytic activity, cleaving dextran on the timescale of crystallisation (several days, 20°C, pH 5.6, Figure S2). This agrees with the dextran digestion assay, in which the total amount of dextran decreased during the assay (Figure S1). Taken together, these results suggest the aspartate nucleophile is more important for catalysis than the glutamate acid/base. The double mutant (GH^dex*^) was also subjected to crystallisation screens, and needle-like crystals formed in similar conditions (Materials & Methods). However, sufficiently large crystals for data collection were not obtained after optimisation.

The entrance to the GH^dex^ binding cavity is stabilised by surface-exposed aromatic hydrophobic residues likely to attract glycans. Aromatic residues including Trp and Tyr are highly enriched in carbohydrate binding sites, with glycan C-H groups interacting with protein π electrons (π-stacking) [23]. Phenylalanine also π-stacks with glycans, but there is no significant enrichment of this residue in glycan binding sites in the Uniprot database [24]. GH^dex^ aromatics around the dextran ligand include Trp260, Tyr221/Tyr437, and His173/His175/His254 which are all ∼5 Å from the resolved dextran. All these residues are on solvent-exposed loops in the catalytic domain of the protein, except for Tyr221, which is on the third β-strand of the (β/α)_8_-barrel domain (β3 in Figure 2). There is little structural change once dextran is bound, with both apo GH^dex^ WT and E360A matching liganded GH^dex^ E360A with Cα RMSDs of ∼0.4 Å. The most notable difference is Asp171, which in the dextran-bound structure bends towards subsite −2 and is likely an important stabilising residue, forming multiple hydrogen bonds to the glycan (Figure 2C). Asp171 and His254, which hydrogen bond to subsite glucoses −2 and −3 respectively in GH^dex^, are equivalent to Asp185 and His268 in *Tp*^dex^, which form similar bonds. Asp171 and Tyr221 are highly likely to be interacting residues, based on their hydrogen bonds with subsite −2 of the resolved dextran (Figure 2). Asp171 is on the loop after β-strand 2, and Tyr221 is on β-strand 3. Tyr437, which points to the glycosidic bond between subsites −3 and −4, is on loop between β-strand 8 and α-helix 8 of the catalytic domain.

### Structure determination of SGBP^dex^

The original construct designed for SGBP^dex^ did not crystallise despite extensive screening. Therefore, a truncated construct was designed to remove the N-terminal Ig-like domain (SGBP^dex^ Δ1-147) which was not predicted to bind dextran based on comparison with the canonical SusF structure [25]. The truncation was based on Alphafold2 [15] models of SGBP^dex^ and SusF, which has a homologous domain architecture to SGBP^dex^, with an N-terminal Ig-like domain followed by three Ig-like domains in tandem that each contain one glycan binding site [25]. These domains were denoted carbohydrate-binding modules (CBM) [25], as they are ∼100 residue β-sheet-rich domains that bound starch without enzymatic activity, although they do not belong to a known CBM family. We have also used this terminology for SGBP^dex^ due to its similar fold and role in PUL48. SGBP^dex^ Δ1-147, containing all three putative CBMs, was successfully crystallised and the structure is highly similar to the Alphafold2 model, albeit with a different orientation between CBM1 and CBM2 (PDB: 9SJI, Figure S3). Crystal soaking and co-crystallisation experiments aiming to bind dextran ligands to SGBP^dex^ Δ1-147 were unsuccessful. For SusF, the CBM3 provides much of the starch-binding activity [25], and mutagenesis of predicted tryptophan residues in SGBP^dex^ CBM3 showed this may also be the case in the dextran-binding protein (Table S2). A cis peptide bond identified in the crystal structure may be important for conformational change in SGBP^dex^ (see Figure S4 and Supplementary discussion).

### Structure determination of SusD^dex^ in the absence and presence of substrate

SusD^dex^ was crystallised and structures were solved to resolutions of 1.65 Å (apo, 9SJK) and 1.7 Å (dextran 1.5 soaked, 9SJJ). As expected, the fold of SusD^dex^ was typically SusD-like, and was highly similar to the Alphafold2 model (Cα RMSD = 0.8 Å). SusD^dex^ contains four tetratricopeptide (TPR) domains which form a right-handed superhelix [26], which constitute the cell surface-facing side (Figure 3A). The ligand binding region (residues 248-381), which is relatively unstructured, is sandwiched between TPR 3 and 4, and forms the periplasmic-facing side. This allows SusD^dex^ to deliver bound ligands to the SusC^dex^ after lid closure (Figure 3B), a mechanism which has been well-established across several homologues [8–10,27,28]. The ligand binding site with bound isomaltopentaose (IMO5; isomalto-oligosaccharide 5-mer) is shown in Figure 3C-D. In the ligand binding region, α9 (293-297) and its surrounding loops provide dextran-binding aromatic residues including Trp284, Trp311 and Tyr103. The α3-containing loop and α4 of TPR1 including Asp82, Glu85 and Arg106 provide a lower surface with an extensive hydrogen bonding network to dextran hydroxyl groups. Asn57 and Arg62 from the first 3_10_ helix and Arg300 from the ligand binding region also interface with the IMO5.

**Figure 3.**
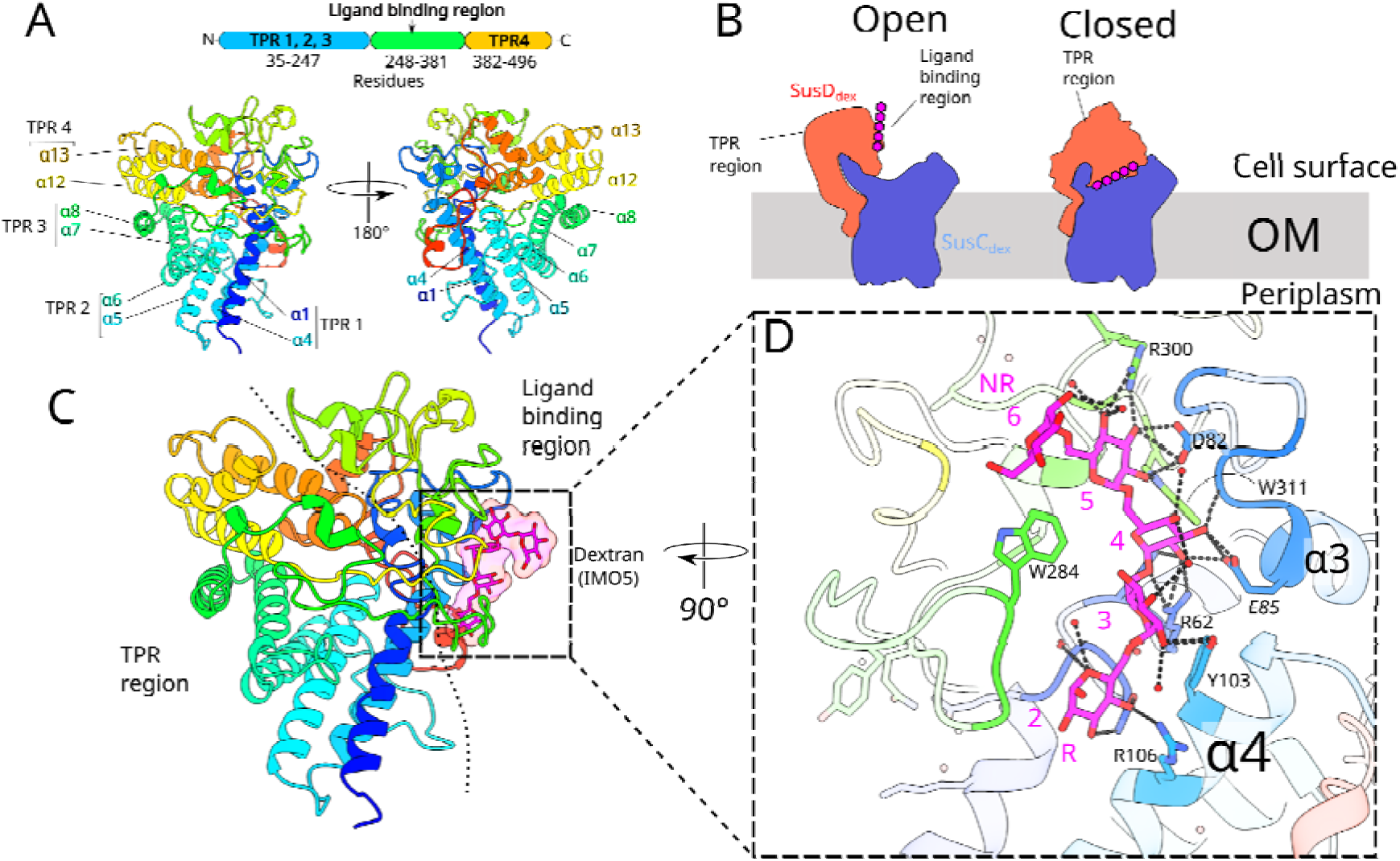
The crystal structure of apo SusD^dex^ and soaked with dextran 1.5. **A**. Apo SusD^dex^ viewed from an arbitrary front and back view. α helices that constitute the four TPR pairs are labelled (α1/4, α5/6, α7/8, and α12/13). In panel **B**, the core utilisome components of the utilisome (SusCD^dex^) are shown in the open and closed states to illustrate the substrate-capturing role of SusD^dex^ in the OM. **C**, Dextran-bound SusD^dex^ with a line separating the two approximate regions of the protein; the more structured and conserved TPR region, and the less structured and conserved ligand binding region. Dextran 1.5 has a DP range between 6-10 residues. The five modelled residues (isomaltopentaose, IMO5) of dextran 1.5 are shown in both stick and surface representation. In **D**, hydrogen bonds with the IMO5 are shown as dashed lines, and IMO5 residues are numbered from the reducing end (R) to the non-reducing end (NR). The numbering corresponds to the subsites defined in the cryo-EM structures (Figure 5). SusD^dex^ structures with and without dextran were deposited under codes 9SJJ and 9SJK, respectively.

Like in GH^dex^, the binding of dextran to SusD^dex^ causes little to no structural change (Cα RMSD of 0.4 Å). Residues along the groove of the binding side including Asp82, Tyr103, Arg106, Trp284, Arg300 and Trp311 have similar side chain conformations between apo and dextran-bound models. The only notable difference is Glu85, which points inwards in the liganded structure, hydrogen bonding with a glycan hydroxyl in subsite 3 (Figure 3D). In the apo structure, the position of the Glu85 carboxylate is instead occupied by a sulphate ion and ordered water, and the residue points away from the binding site.

For comparison between *B. theta* SusDs, surfaces (coloured by hydrophobicity) of liganded SusD^dex^, SusD^starch^ and SusD^lev^ are shown in Figure S5. Cα RMSDs of SusD^dex^ when matched to SusD^lev^ and SusD^starch^ were 6.9 Å and 9.3 Å respectively. The SusD^dex^ dextran binding site has a solvent-exposed Trp284, below which a hydrophobic groove in the protein surface is formed by Trp292, Trp311, Phe296, Trp309 and Tyr370. These residues stem from the ligand-binding region (Figure 3C). Across from Trp284, the N-terminal α-helix 4 (which forms TPR 1 with α1) provides a solvent-exposed Tyr103 which appears to clamp the ligand between Trp284 and Tyr103. The shape of the SusD^starch^ binding site appears to be divergent from the levan and dextran SusDs (Figure S5). Unlike SusD^dex/lev^, there is no central protruding aromatic residue (Trp284 and Phe649 for SusD^dex^ and SusC^lev^ respectively) and the ligand projects from the protein surface, instead of running parallel like for SusD^dex/lev^. This may be related to the smaller helical pitch of starch, resulting in a more curved substrate [29]. The groove of the binding site is more pronounced, with aromatic residues Trp85 and Tyr296 protruding from the surface, with the substrate in between. Trp96 and Trp320 form an aromatic patch between the protruding residues. The SusCD^lev^ site more closely resembles the SusD^dex^ site (Figure S5C). In SusCD^lev^, the solvent-exposed Phe649 of SusC^lev^ provides the central aromatic group, equivalent to Trp284 in SusD^dex^. Below Phe649 is a groove of hydrophobic and aromatic residues of SusD^lev^ including Phe301 and Tyr395. Like Tyr103 and Trp284 in SusD^dex^, Trp85 (SusD^lev^) and Phe649 (SusC^lev^) may provide a clamp to hold levan substrate, although only Trp85 is essential for levan binding [9]. Trp85 may be replicated in SusD^dex^ by Tyr103, which occupies a similar position in α4 of TPR 1 in both SusDs.

### Cryo-EM of the inactive dextran utilisome in the presence of ligand

We previously established that dextran and levan utilisomes are dimeric four-component complexes that are stable in the absence of substrate [10]. To investigate how substrates bind to and potentially affect the dextran utilisome, we first generated a *B. theta* strain with the GH^dex*^ double mutation D297A/E360A and a His6 tag on the C-terminus of SusD^dex^ to allow the purification of the catalytically inactive utilisome. Several approaches were tried to obtain a sample that behaved well on cryo-EM grids (Methods). The best dataset was achieved through a combination approach. DDM was used for complex extraction and was then exchanged to lauryl maltose neopentyl glycol (LMNG) during IMAC. Excess LMNG was removed by SEC in the absence of detergent (see Figure 4A). For vitrification, UltrAuFoil grids were PEGylated with a protocol to render the gold support layer hydrophilic [30], and a secondary detergent (fluoro-octyl maltoside) was added before vitrification. This led to more abundant particles that were distributed throughout the grid holes. A mixture of dextran 1.5 and 3 was added at 0.5 mM each and incubated for 30 mins before vitrification. Following data processing (Figure S6, S7, Table S3), the resulting reconstructions had nominal resolutions of ∼3 Å based on gold-standard Fourier shell correlation of the masked volumes (Figure S7), with density for all protein chains of the octameric complex (2 x SusC^dex^, 2 x SusD^dex^, 2 x GH^dex^ and 2 x SGBP^dex^). The SGBP^dex^ density remained low-resolution, likely due to the flexibility of the protein, and only the N-terminal domain (NTD) could be modelled (Figure 4).

**Figure 4.**
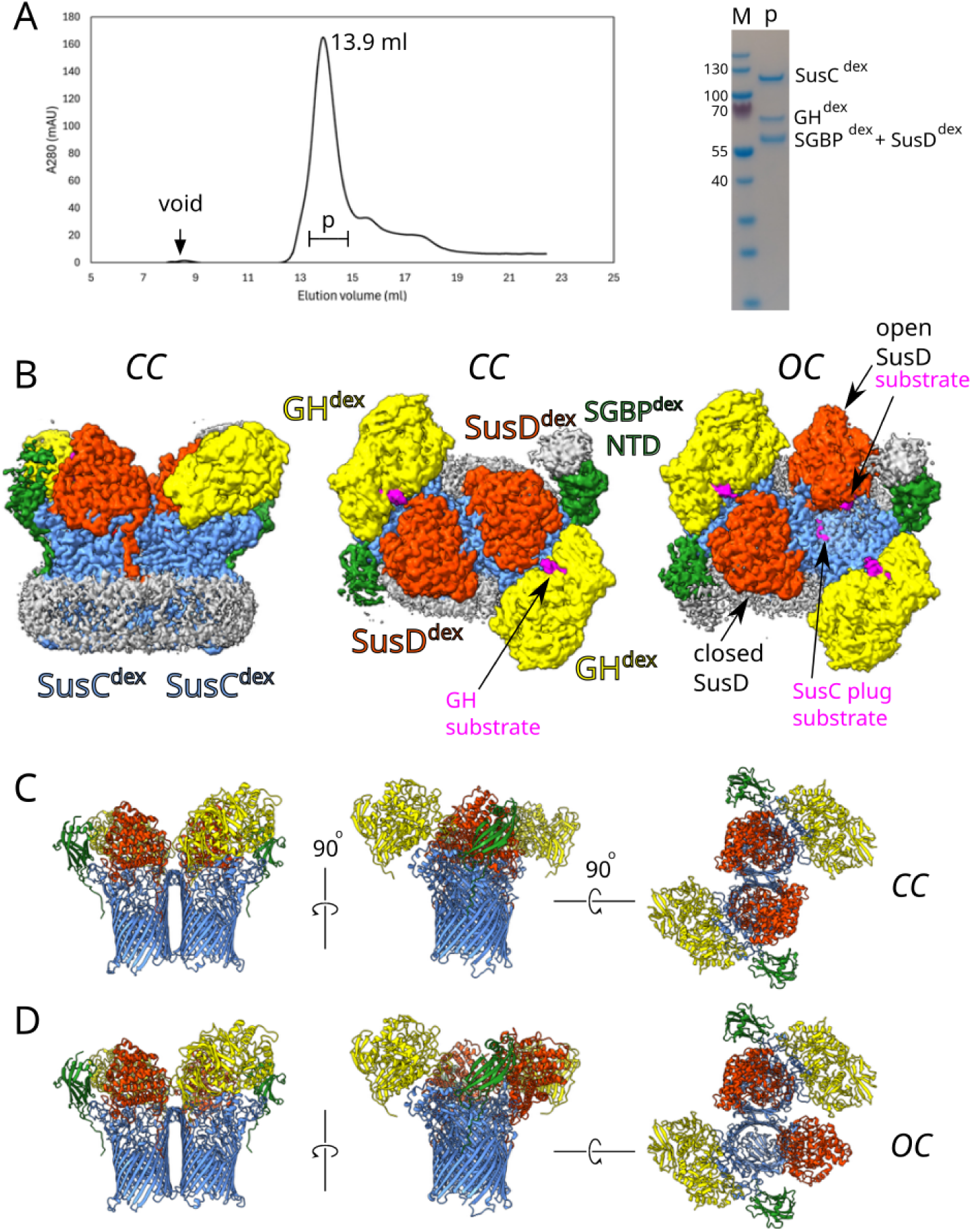
The structure of the inactive dextran utilisome bound to ligand. **A**. Superose 6 SEC profile and SDS-PAGE gel of the purified dextran utilisome. “p” denotes the SEC peak fractions that were pooled. **B**. Final 3D reconstructions of the closed-closed (CC) and the open-closed (OC) states of the inactive dextran utilisome, in complex with dextran. Closed-closed and open-closed maps had nominal resolutions of 2.5 and 2.9 Å (Figure S7C), after FSC-mask auto-tightening, respectively. The map was colour zoned according to the atomic models of the dimeric four-component complex (GH^dex^:SGBP^dex^:SusD^dex^:SusC^dex^); GH^dex^ is yellow, SGBP^dex^ is green, SusD^dex^ is orange-red and SusC^dex^ is cornflower blue. Densities with modelled isomaltooligosaccharides are magenta. Note the additional substrate densities visible in the open state. Only the N-terminal domain (NTD) of SGBP^dex^ was reconstructed at a modellable resolution, but the putative three carbohydrate-binding modules (CBM) are visible at higher thresholds (Figure S9). **C, D** Cartoons of the modelled dextran utilisome in various orientations for the CC (**C,** 9SM2) and OC (**D,** 9SJE) state.

### The dextran utilisome remains dynamic in the presence of substrate

Previous work on the active dextran utilisome in the absence of substrate found only classes with open SusD lids after 3D classification [10]. In the inactive dextran utilisome plus substrate reported here, ∼60% of particles had closed SusD^dex^ lids (“closed-closed”, CC, 9SM2), while ∼40% were observed with one SusD open (“open-closed”, OC, 9SJE) (see Figure 4 and Figure S7A). This differed from the inactive levan utilisome where only closed states were observed upon substrate addition [10]. Additionally, SGBP^lev^ was tethered to the rest of the utilisome by fructose oligosaccharides (FOS) of DP 15-25, which enabled its model building. By contrast, dextrans between 1 and 20 kDa did not have the same effect on SGBP^dex^ across several grids and datasets, suggesting a subtly different role for SGBP^dex^.

The dynamics within the dextran utilisome were visualised via 3D variability analysis [31], showing the closing of SusD^dex^ (Movie S1 and Figure S8). At the beginning of the movie (Figure S8, Frame 0), the C-terminal domain (CTD) of SGBP^dex^ contacts the active site of GH^dex^, which may be how SGBP^dex^ supplies dextran to GH^dex^ for hydrolysis. As SusD^dex^ hinges closed, SGBP^dex^ becomes poorly resolved, but at the end of the movie the SGBP^dex^ CTD can be seen above the closed SusD^dex^ (Frame 59), translating to an approximate 40 Å shift in position of the C-terminus of the protein. To estimate the positions of SGBP^dex^ in low contour density, Alphafold-predicted domains CBM 1 and CBM2-3 were docked independently in the CC and OC maps (Figure S9). Docking into the OC map showed that the density is weakest at CBM2, indicating that this domain is the most mobile. Cis/trans isomerisation of a cis-peptide in CBM2, Ala306-Asp307 (Figure S4), may help drive movement of SGBP^dex^ CBM domains upon dextran binding (see Supplementary discussion, and Figures S4, S8-9).

### Two extracellular hinge loops of SusC^dex^ are involved in the closing of SusD^dex^

In the dextran utilisome, extracellular loops 7 and 8 of SusC^dex^ act as a mobile platform for SusD^dex^, similar to the levan utilisome [10]. Here, to emphasise their importance, SusC^dex^ loops 7 and 8 are termed “extracellular hinge loops” (L7, EHL1: 627–667 and L8, EHL2: 697–722) (Figure S10). Movement of EHL1 and EHL2 (Figure S10B-D) and polar interactions with SusD^dex^ (Figure S11) may be implicated in stabilising the closure of substrate-loaded SusD^dex^. PDBePISA [32] analysis was performed, with SusC-D^dex^ interfaces and polar interactions in the open and closed states included in the supplementary information (Spreadsheet S1). EHL1 and EHL2 make up 48% and 64% of SusC^dex^ polar interactions with closed and open SusD^dex^, respectively. SusC^lev^ EHL1 is longer than the SusC^dex^ EHL1 by approximately 20 residues. In the closed state, these residues project towards captured ligand, providing a binding surface with SusC^lev^ Phe649 at the tip of the loop (Figure S5C) [10]. Other important loops are loop 1 (253–303) and loop 9 (759–843) which act as binding platforms for GH^dex^ and SGBP^dex^, respectively. There is almost no conformational change in loops 1 and 9 between the open and closed models, suggesting that they are static.

The open and closed dextran utilisome cryo-EM models were analysed, revealing an opening of 62°, with a small shift of 2 Å (Figure S10A). This was a relatively large opening compared to the open and closed models from the core levan utilisome ([SusCD]_2_), which only had a measured rotation angle of 23° (1 Å shift). A state of the apo levan utilisome with a larger opening was previously observed, but it was not modelled [9,10]. The fully-open SusCD^dex^ opening is similar to open states of both ButCD and RagAB (from *Bacteroides fragilis* and *Porphyromonas gingivalis*, respectively [27,28]) based on visual inspection (Figure S12).

Between the open and closed SusC^dex^, the tip of EHL1 shifts 21.7 Å inwards during lid closure (based on Cα positions of Gly647). Thr708, at the tip of EHL2, undergoes a smaller shift of 10.6 Å (Figure S10C-D). Based on the loop structures in both states, EHL1 moves as a rigid body akin to a lever, while EHL2 undergoes a rotation towards the SusC^dex^ lumen stemming from the 2-stranded β-sheet at the base of the loop. The phenyl ring of EHL2 Phe718, at the top end of the β-sheet, is shifted 7 Å towards EHL1 after lid closure (based on the phenyl ring *p*-carbon). Another consequence of the rotation is a shift of SusC^dex^ Asp713 Cα of 6.5 Å inwards towards SusD^dex^ Arg106 in the closed state, forming two salt bridges (Figure S10E). Arg106 itself moves ∼16 Å inwards during closure, indicating these interchain salt bridges may stabilise the closed state during the transport cycle.

EHL1 forms 14 and 26 close contacts (Van der Waals overlap > −0.1 Å) with SusD^dex^ in the open and closed states, respectively. A contact in the open state to SusD^dex^ Pro166 is not maintained during closure, although nearby Phe164 contacts EHL1 in both states. Most of the additional contacts in the closed state come from α1 of SusD^dex^ (residues 39-54). A hydrophobic motif (Tyr644-Pro645-Phe646) at the tip of EHL1 contacts SusD^dex^ α1, particularly Gln54, and loop residues at the beginning of the unstructured ligand binding region (Gln274, Gln289). SusC^dex^ Tyr644 and SusD^dex^ Tyr50 appear to form an aromatic stack which is maintained during closure. After closure, residues on the loop between TPR 2 and 3 contact EHL1, including Arg208 and Asn204. By contrast, EHL2 makes 4 and 8 contacts to SusD^dex^ in its open and closed states, respectively. In the open state, three of the four contacts are to SusD^dex^ Phe164, which are broken when SusD^dex^ closes, where Phe164 contacts the closed state EHL1 instead. This suggests that SusD^dex^ Phe164 is an important residue for the hinging movement. After EHL2-Phe164 contacts are lost during closure, two tryptophans in α4 of SusD^dex^ TPR1, Trp105 and Trp109, form an interface with EHL2.

In the closed conformation, there is one close contact between SusD^dex^ Arg348 and GH^dex^ Asp364, which is not observed in the open conformation. Closer inspection revealed that an electropositive patch of SusD^dex^ (Arg348, Arg347) is near an electronegative patch of GH^dex^ (Glu365, Asp364, Asp367), which may provide rigidity in the closed complex. These residues do not form salt bridges in the model, but the inter-residue distances appear to be in the proper range for interaction (3-4 Å). The levan utilisome does not share this feature. The modelled portion of SGBP^dex^ (N-terminal domain; 21-154) forms no contact with SusD^dex^ or GH^dex^ in either the open or closed state.

### Substrate binding sites in the dextran utilisome

The binding sites of GH^dex^, SusD^dex^ and SusC^dex^ in the open and closed utilisome models are shown in Figure 5 and Movie S2 (plus Table S6), with an overview in Figure S13. For GH^dex^, differences between the open and closed states are very minor with a Cα RMSD of 0.5 Å. GH^dex^ has numerous aromatic glycan binding residues that define the IMO7 (OC) and the IMO6 (CC) binding pocket (Figure 5A-B). From the “top” (OC, subsite −5) to the “bottom” (subsite +2) these are; Trp260, Tyr437, His254, His176, Phe438, Tyr221, Tyr396, Tyr393, and Trp362. When viewing GH^dex^ from the direction of SusD^dex^, three residues (Trp362, Tyr393 and Tyr396) form the right wall of the binding site. In the closed state, these three residues are shifted slightly inwards, away from SusD^dex^ that comes into close contact with GH^dex^ after hinging. Trp260, at the top of the binding site, is important for dextran binding given that its substitution by Ala led to a 62-fold decrease in K_d_ for dextran 1.5 (DP = 6-10) (Table S2). A substitution of Trp362 to Ala could not be expressed, indicating that it may be required for protein folding. At these resolutions (> 2.0 Å), it is not possible to confidently say which is the correct orientation of the glycan chain. The reducing end was chosen to point downwards, towards SusD^dex^ in both models (Figure 5A, B). In the closed GH^dex^ model, the IMO6 conformation is similar to the crystal structure of Tp^dex^ [20] (Figure S2D), with the non-reducing end (subsite −4) pointing away from SusC^dex^ (Figure 5B), and from the catalytic residues at the bottom of the binding site.

**Figure 5.**
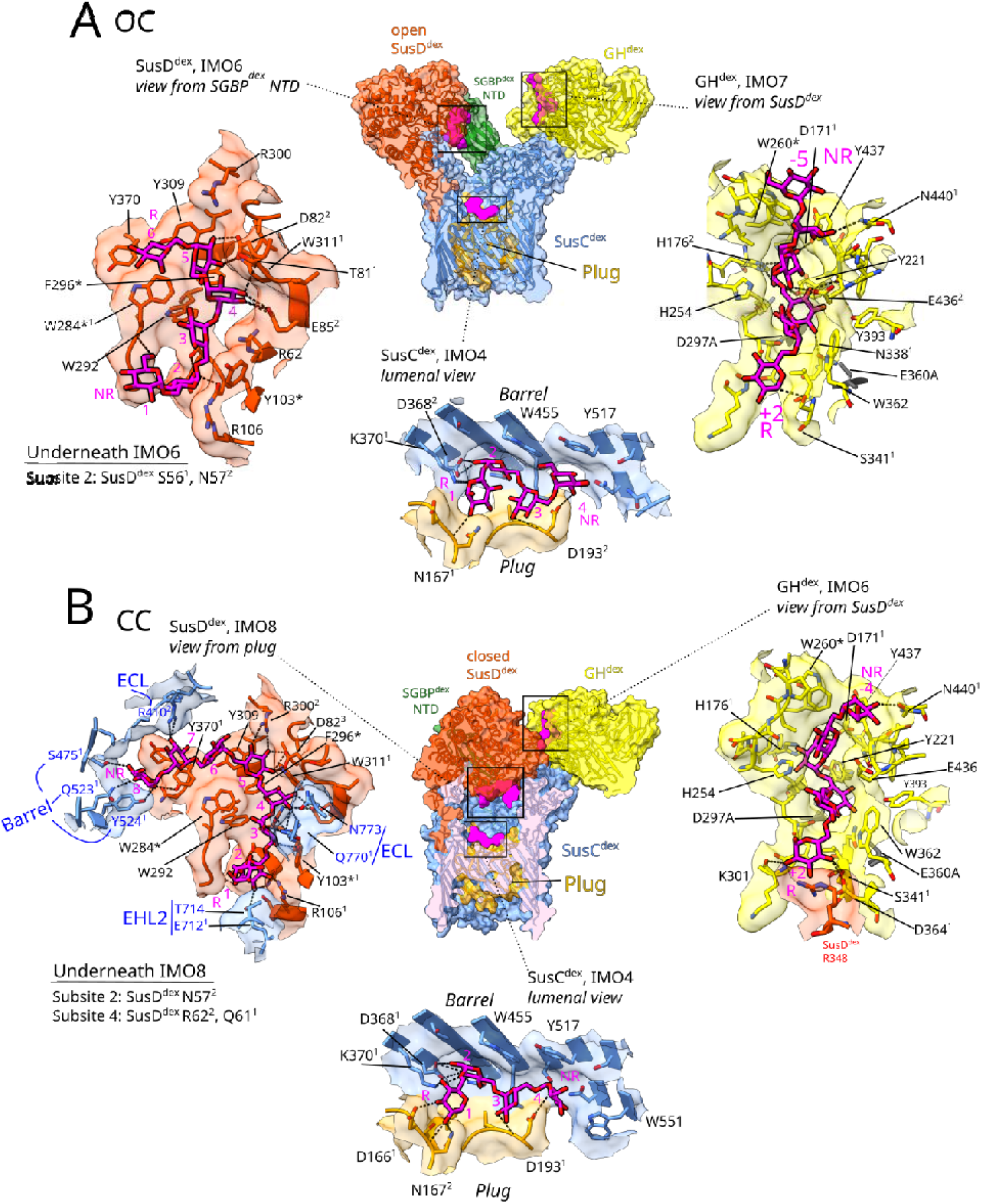
Dextran binding sites in GH^dex^ (yellow), SusD^dex^ (orange-red) and SusC^dex^ (blue) in the open (A) and closed (B) utilisome. Colour scheming is consistent with previous figures, with the models are clipped to view the lumen of SusC^dex^. Overviews of the utilisomes are shown in the centre. Dextran oligosaccharides are shown in magenta, with protein-glycan hydrogen bonds shown as black dotted lines. “IMOx” are isomalto-oligosaccharides where x denotes the number of α-LJ-glucose residues. Superscript numbers on the residues denote how many atoms are hydrogen bonding to dextran. For both models, each binding site is magnified and displayed at the most informative orientation. Protein residues up to 5 Å from the ligands are displayed in stick representation, with aromatic and other notable residues are labelled. The subsites are numbered by glucose residue, with non-reducing (NR) and reducing ends (R) labelled. For GH^dex^, these follow the convention of increasingly negative numbering from the cleavage site towards the non-reducing end [33]. SusC^dex^ residues are grouped and labelled based on what part of SusC they originate from; EHL2 and ECL refer to extracellular hinging loop 2 and extracellular loops, respectively. Barrel refers to residues on the beta-strands, and plug refers to the SusC^dex^ plug domain (residues 129-232).

Closed state SusD^dex^ has the largest bound dextran ligand observed (IMO8), whereas in the open state only part of the binding site was stably occupied (IMO6, subsites 1-6). Since the local map resolution of the open SusD^dex^ was much lower than its equivalent in the closed state (∼3-5 Å vs ∼2 Å, Figure S14), it is not surprising that less dextran density was observed. The lower resolution is likely due to flexibility of the open SusD^dex^, and the fact that the OC map had many fewer particles than the CC map (∼21,000 vs. 116,000). By contrast, the closed SusD^dex^ state is stabilised by several interactions with SusC^dex^, minimising flexibility (Figure S11).

Like in the crystal structure of liganded SusD^dex^, dextran forms a planar semi-circle around the central upper aromatic binding residue Trp284. Based on ITC, Trp284 and Phe294 were essential for binding of dextran 1.5, while Tyr103 was beneficial but not necessary (Table S2). Closed SusD^dex^ forms 12 contacts with its IMO8 ligand (Van der Waals overlap > −0.01 Å), primarily via hydrogen-bonding residues, including Asp82, which has 3 contacts with subsite 5. Near the reducing end, subsite 2 is also closely associated with SusD^dex^, with 2 contacts with Asn57, and 1 with Arg106. Moving along the glycan, subsite 3 contacts Tyr103, while subsite 4 contacts Gly83. Tyr370 also makes a contact within its phenolic ring to subsite 7. Interestingly, compared to closed SusD, open SusD^dex^ forms more contacts (17 *vs*. 12) with its smaller IMO6 ligand. Contacts with hydrogen bonding residues Asp82, Asn57 and Arg106 are maintained, but additional aromatic-ligand interactions are present, particularly at subsite 6, which has 3 contacts to Trp284 and 2 to Tyr307. While the overall ligand poses between the crystal structure and both cryo-EM states of SusD^dex^ are highly similar, the open SusD^dex^ ligand differs at its terminal subsite 6, which bends inwards towards Trp284. This suggests that Trp284 provides π-stacking interactions to dextran oligosaccharides during transport that may be broken after lid closure, contributing to ligand release.

In the closed state, SusC^dex^ has several residues that hydrogen bond with IMO8. These SusC^dex^-glycan hydrogen bonds (Figure 5B, in blue) and SusCD^dex^ salt bridge interactions (Figure S11E and S15) likely help stabilise the closed state. Starting at the non-reducing end, several residues at the top of adjacent β-barrel strands bond to subsite 8 (Ser475, Gln523, Tyr524, Figure 5B). Extracellular loop (ECL) residues Arg410 and Gln770 bond to subsites 7 and 3, respectively. EHL2 Glu712 hydrogen bonds to subsite 2, while Thr714 of EHL2 forms a close contact to subsite 1, which is the only SusC^dex^ residue to do so, possibly indicating subsite 1 as a favoured endpoint for substrate loading (Figure 5B, EHL2). Additionally, Asp713 is sandwiched between these residues, and forms a protein-protein salt bridge with the nearby SusD^dex^ R106 (Figure S11). This suggests the EHL2 ^712^Glu-Asp-Thr^714^ motif may have a multi-functional role in both ligand stabilisation and lid closure.

When the contour of the CC map is lowered (Figure S15, contour 0.026), the closed SusD^dex^ IMO8 density extends further another two glucose residues towards the binding site of the SusC^dex^ plug domain, but this section of density was not modelled. SusC^dex^ Tyr477 is proximal to the unmodelled density at IMO8 (Figure S15, model only), which like Tyr542, faces the interior of the barrel. At still lower map contour (contour 0.017-0.010), the IMO density extends from both ends of IMO8 (A and B) towards the SusC^dex^ plug site, while IMO4 (C and D) extends in the opposite direction, suggesting some complexes contain longer dextran oligosaccharides that span both the SusD^dex^ and SusC^dex^ binding sites simultaneously. Based on this weak density, we estimate a dextran size limit of around DP 17 (∼3 kDa) could be present in the SusCD^dex^ cavity, in rough agreement with the levan utilisome [10].

In the SusC^dex^ plug binding site, isomaltotetraose (IMO4) was modelled in both states, although the density was weaker for open SusC^dex^. In both models, the main glycan binding residue appears to be SusC^dex^ Trp455, from the luminal face of β-strand 7. SusC^dex^ conformations around both open and closed plug sites are almost identical, indicating no major changes occur at the plug between open and closed states. Subsites 1 and 2 form the majority of close contacts to Lys370, Gly425 and Trp455 of the β-barrel, as well as to Asp166 and Asn167 of the plug domain. Residues involved in hydrogen bonding include β-barrel residues Asp368 and Lys370, as well as Asp166/193 and Asn167 from the plug domain (Figure 5A and B, orange surface). IMO4 conformations are highly similar for subsites 1 and 2. In the open transporter, the IMO4 subsites 3 and 4 are bent inwards, with the subsite 4 π-stacking with Tyr517, while in the closed state, IMO4 subsite 4 is wedged between Tyr517 and Trp551. Although the side chain of Trp551 is not displayed in Figure 5A for the open state (due to the different IMO4 conformation and the 5 Å radius requirement from the ligand for residue display), it is in an almost identical conformation to that in the closed state (Figure 5B).

### The dextran utilisome, like the levan utilisome, has an aromatic lock

The “aromatic lock” of SusC-type TBDTs was first described in the levan utilisome [10]. Comparison of the substrate-bound and apo levan utilisome structures showed that in the apo utilisome, Tyr89, Tyr191 and Phe558 form a triple aromatic stack that physically links the TonB box (via Tyr89, which is next to TonB box residues Asp82-Gly88), the plug domain (via Tyr191) and the inner wall of the β-barrel (via Phe558). Upon substrate binding, Trp685 shifts further into the barrel, moving Phe583 up and Ser193 down. This movement causes the Tyr191-containing-loop to move further towards the periplasm, breaking its interaction with Tyr89 and releasing the TonB box-containing loop into the periplasm. Consequently, in the substrate-bound structure this loop is not modelled as it is too flexible to be resolved, but the last resolved residue in both structures (Arg93) undergoes a 35 Å shift. This displacement/disordering of the SusC N-terminus is likely necessary for signalling substrate binding across the OM, allowing the TonB protein to access the TonB box sequence in the periplasmic space (Figure S16).

The aromatic lock is conserved between the levan and dextran utilisomes. It is important to note that residue numberings are shifted between these structures, as the levan utilisome polypeptides were numbered from the first mature residue whereas the dextran utilisome polypeptides were numbered including their N-terminal signal sequences (*i.e*. the SusC^lev^ numbering is offset by −25). The SusC^dex^ β-barrel-loop residues Phe582 and Trp666 are homologous to SusC^lev^ Phe583 and Trp685. SusC^lev^ Phe558, which is internally facing from a β-strand, is absent in SusC^dex^, with a methionine in its place (Met557). This means the aromatic stack in the dextran utilisome is a “double” (Tyr219 and Tyr117) instead of a “triple” stack described for the levan utilisome [10]. However, methionine residues are a common bridge between two aromatic residues in an Aro-Met-Aro fashion [34], indicating that Met557 likely has a similar role to SusC^lev^ Phe558, as it occupies the same position on the barrel wall (Figure S16D).

Allosteric substrate signaling appears to begin at SusC^dex^ Trp666, at the end of EHL1, where it flips towards the plug in the closed state (Figure S16). This has two consequences. Firstly, Phe582 is nudged upwards by Trp666, slightly reordering the β-barrel-loop residues 579-587. This causes Lys584 to point towards SusD^dex^ Asp24 on the N-terminal linker of the the SusD^dex^ lid, forming an additional salt bridge (to Asp24-Arg587, which is present in both open and closed states, Figure S11) which may stabilise lid closure (Figure S11, S16B-C). A similar trend is observed in the position of equivalent lysine Lys585 in the active (open) and substrate bound (closed) levan utilisome structures, with Lys585 forming a salt bridge with SusD^lev^ Asp6 in the closed state (Figure S16D). Secondly, the downwards movement of the Trp666 indole ring causes Ser221 to flip away, carrying the signal to the plug domain. Ile218 and Tyr219 clash with Tyr117 of the N-terminal loop, breaking the Tyr-Met-Tyr stacking interaction between Tyr117, Met557 and Tyr221. The resulting N-terminal disordering and movement allows the TonB box to be accessed by TonB in the periplasmic space: the first residue resolved in closed SusC^dex^, Thr119, is shifted ∼16 Å slightly down and away to the opposite end of the barrel lumen compared to its position in the open state (Figure S17). The first resolved residue of open SusC^dex^ (Leu109) occupies a similar position to where Thr119 is in the closed model.

### Isothermal titration calorimetry shows varying affinities of the SLPs for dextrans

Dextran-binding activity was quantified by titrating dextrans from 1.5 to 500 kDa into purified SLPs (Table S4). SusD^dex^, SGBP^dex^, SGBP^dex^ Δ1-147, and two catalytically inactive mutants of GH^dex^ (E360A and D297A/E360A) were used. GH^dex^ had the highest binding affinity for dextran (1-4 µM), independent of dextran size. SusD^dex^ affinities were lower, in the 8-20 µM range. Both GH^dex^ and SusD^dex^ showed little difference between the size of dextran and K_D_. By contrast, dextran binding affinities of SGBP^dex^ and SGBP^dex^ Δ1-147 depended strongly on ligand size (Figure 6).

**Figure 6.**
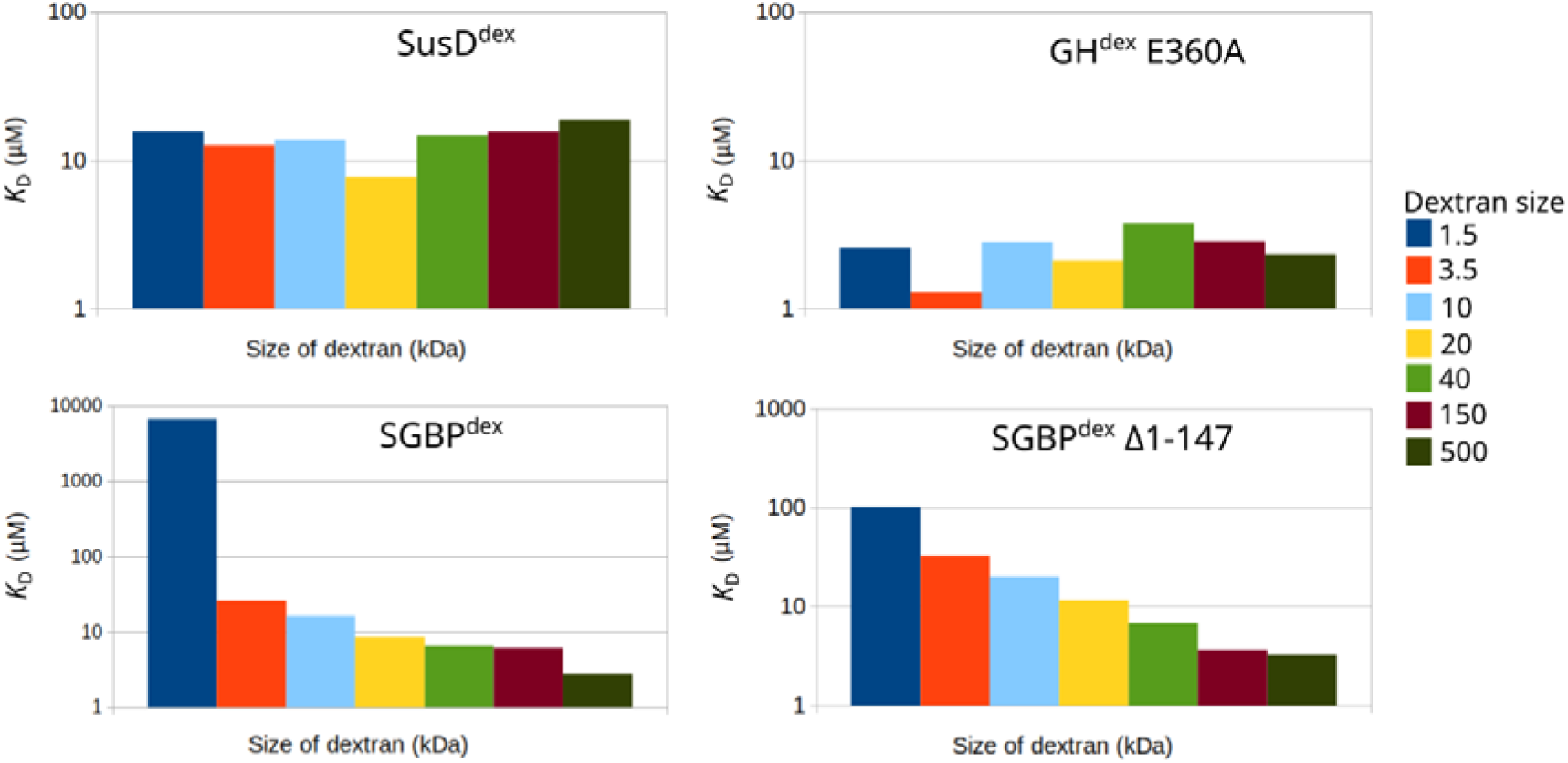
Substrate affinities (µM) of SusD^dex^, GH^dex^ E360A, SGBP^dex^ and SGBP^dex^ Δ1-147 against dextrans from 1.5 to 500 kDa. Derived K_D_ values from Table S4 are plotted on a logarithmic scale against dextran size. Bars are colour-coded by dextran size. Note that the Y axes differ in scale between the different proteins. Example titrations are shown in Spreadsheet S2.

SGBP^dex^ had a very low affinity (6 mM) for dextran 1.5 with high variability between measurements due to the poor suitability of ITC to measure low-affinity interactions. Affinities increased with the size of dextran; when SGBP^dex^ and SGBP^dex^ Δ1-147 were titrated with dextran 500, K_D_ values comparable to GH^dex^ were obtained (∼3 µM), indicating that SGBP^dex^ preferentially binds large dextrans that have not yet been processed by GH^dex^. The number of binding sites of the three SLPs were also analysed using ITC (see Table S5 and Supplementary discussion).

## Discussion

Figure 7 shows the proposed transport cycle of the dextran utilisome, which was based on both the cryoEM-derived OC and CC models, in combination with the 3D variability data. In step 1, CBM 3 of SGBP^dex^ contacts the GH^dex^ active site (Figure S8-9), supplying dextran to be hydrolysed endolytically [16]. The products then bind to the open SusD^dex^ lid. The observation of bound ligand to open SusD is interesting and suggests that ligand binding does not immediately result in lid closure akin to a venus flytrap mechanism (at least *in vitro*). Some dextran is likely released into the environment post-GH^dex^ hydrolysis, facilitating the cross-feeding of other microbiota members [35]. When the SusD^dex^ lid closes the transporter lumen is loaded (step 2). An open question is whether lid closure occurs spontaneously (due to natural dynamics of the lid) or due to a yet unknown trigger. Previous single-channel physiology experiments on plug-less SusCD^lev^ in the absence of substrate showed very noisy current traces with large amplitudes (∼1.5 nS), indicative of rapid (< ms) opening and closing of the SusD lid [8]. In addition, cryo-EM structures of apo SusCD^lev^ lacking the additional SLP components show open and closed SusD lids [9]. A prerequisite for the spontaneous dynamics model is that closure of an empty SusD lid should not lead to TonB box exposure since that could result in a non-productive transport cycle. However, complete levan utilisomes purified under mild conditions showed exclusively open SusD lids without substrate and closed lids with substrate [10], suggesting a substrate binding-driven lid closure *in vivo*. Indeed, closed SusCD complexes *without* bound ligand have not yet been observed, and we therefore favour a substrate-driven model of “stable” lid closure. This model might explain why TBDT lids evolved in the first place: if it takes a long time for the interaction with TonB to occur, a stably closed lid would prevent dissociation of the ligand back into the extracellular space, as would happen with a regular, “lidless” TBDT. Another consideration is that it is not clear what influence, if any, the native OM environment (likely dominated by negatively charged lipo-oligosaccharides on the cell surface [36]) has on lid dynamics.

**Figure 7.**
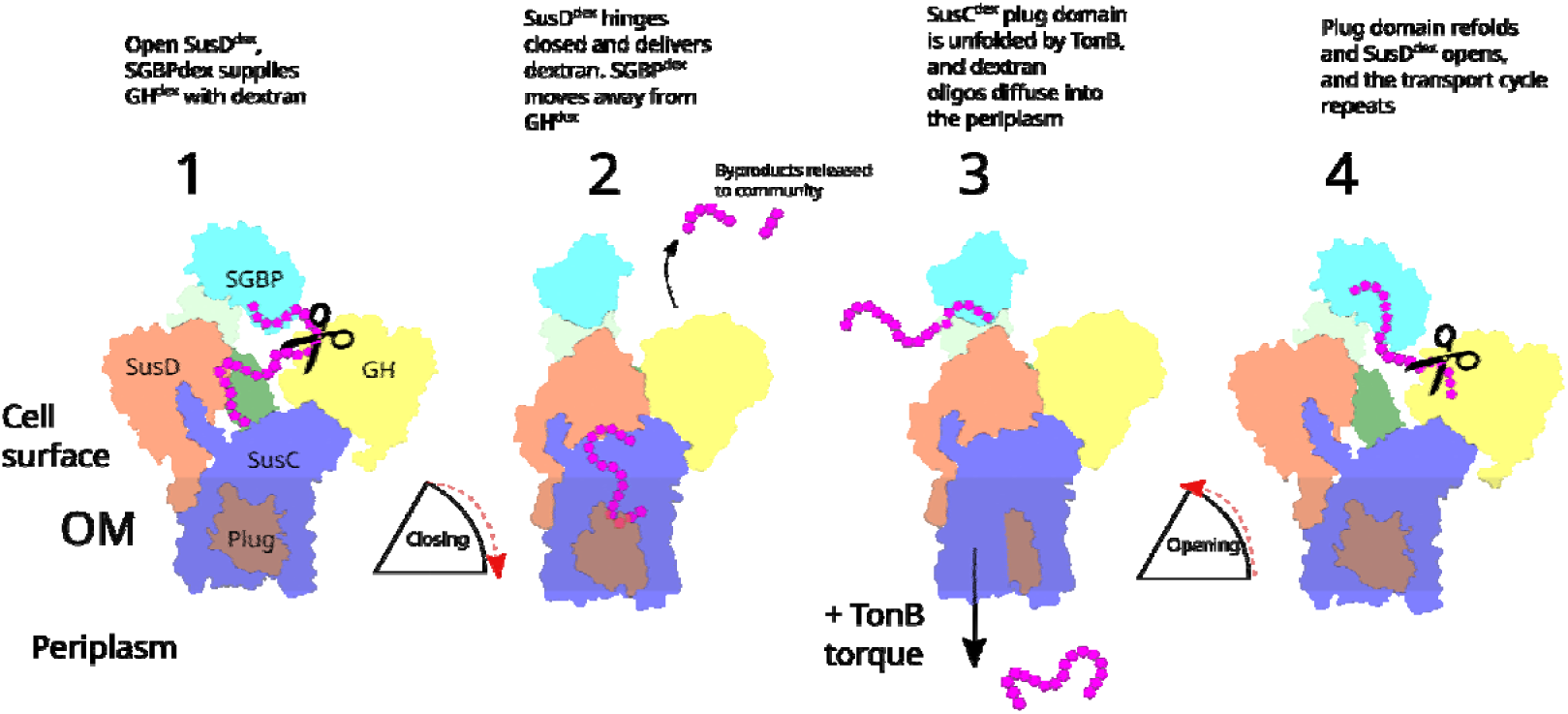
A schematic representation of the dextran utilisome transport cycle. Four-component utilisomes, rather than the full octameric complexes, are shown for clarity. Side views based on the OC (1 and 4) and CC (2 and 3) models are shown in the OM plane. GH^dex^ is yellow, SusD^dex^ is orange red and SusC^dex^ is cornflower blue. For SGBP^dex^, the dark green portion is the modelled NTD, while the pale green and cyan sections show the docked positions of CBM 1 and CBM 2 + 3, respectively. Magenta pentagons are dextran oligosaccharides.

During lid closure, SGBP^dex^ moves away from GH^dex^ (Figure S8), and stabilising salt bridges are formed between SusC^dex^ and SusD^dex^ (Figure S11, Supplementary information). An allosteric signal cascade travels down SusC^dex^ to disrupt the aromatic lock, causing a conformational change of the SusC^dex^ N-terminus that exposes the TonB box for interaction with the inner membrane TonB-ExbBD complex [37,38] (step 3; TonB-ExbBD complex not shown). Rotary proton pumping by ExbBD causes conformational differences in TonB, generating a pulling force which then partly unfolds the SusC^dex^ plug domain, creating a transient pore [39]. IMO products are then able to diffuse into the periplasm. After dissociation of the TonB box-TonB interaction, the plug refolds and SusD^dex^ opens for a new transport cycle (step 4).

## Materials and Methods

### Cloning of SusD^dex^, SGBP^dex^ and GH^dex^ for *E. coli* expression

The following protocol was used to generate soluble constructs for the PUL48 surface-exposed lipoproteins *BT3087*, *BT3088* and *BT3089.* To facilitate crystallisation, primer pairs were designed for each construct to remove the N-terminal signal sequence and an additional ten amino acids of the mature lipoprotein, including the cysteine that would be tri-acylated in *B. theta*. Genomic sequences of *Bacteroides thetaiotaomicron* VPI-5482 for primer design were obtained from KEGG (https://www.genome.jp/), and primers were synthesised by Eurofins genomics. N-terminal signal sequences were predicted by supplying the amino acid sequence to SignalP 5.0 [40]. The constructs contained residues 26-592, 31-504, and 29-496 for *BT3087*/GH^dex^, *BT3088*/SGBP^dex^, and *BT3089*/SusD^dex^ respectively. *BT3088 (*Δ*1-147)* contained residues 148-504, corresponding to a deletion of the N-terminal domain. Sequences of *BT3087*, *BT3088* and *BT3089* were amplified from purified *B. theta* genomic DNA with primers containing NcoI and XhoI restriction sites, restricted and ligated into pET28b, giving the constructs a C-terminal His6-tag for purification. Chemically competent *E. coli* TOP10 cells were then transformed using heat shock. Positive *E. coli* TOP10 clones were identified by colony PCR of individually picked colonies that were grown on an antibiotic selection plate after ligation. These constructs were designed to be overexpressed in the *E. coli* cytoplasm by IPTG-induction of the *T7lac* promoter.

### Expression of SusD^dex^, SGBP^dex^ and GH^dex^

LB media (Melford Laboratories) was dissolved in deionised water at 20 g/L and autoclaved. BL21 (*DE3*) precultures, inoculated from fresh transformations on LB-agar plates with antibiotic, were first propagated overnight at 37 °C and 1/100^th^ of the second culture volume was typically used as inoculum (*e.g*., 10 ml per 1L final culture). Cells transformed with the pET28b vector were selected using 50 µg/ml kanamycin (Melford). After growth to exponential phase (OD600 = 0.4-0.8), Isopropyl ß-D-1-thiogalactopyranoside (IPTG, Melford) was added to final concentrations between 0.2 mM and 1 mM to induce protein expression. Different expression conditions after induction were tested, such as 37 °C at 180 rpm for 2-4 hours or 16-20°C, 150 rpm for 19-22 hours until acceptable conditions were found for each construct.

### Cloning of *BT3087* (GH^dex^) catalytic mutants and dextran binding mutants in *BT3087*, *BT3088* and *BT3089*

Putative catalytic residues in *BT3087* were identified from the literature [16,17]. The Q5 site-directed mutagenesis kit (New England Biolabs) was used to introduce the D297A and E360A mutations in the previously cloned pET-28 *BT3087* construct using a novel primer pair designed with New England Biolab’s online tool (https://nebasechanger.neb.com/). This protocol involved whole-vector PCR with primers harbouring the gene modifications. After the single mutants were confirmed by Sanger sequencing (Eurofins Genomics), a double mutant (D297A, E360A) was produced using a confirmed single mutant as template DNA. The same method was used to create dextran-binding mutants of *BT3087* (GH^dex^), *BT3088* (SGBP^dex^) and *BT3089* (SusD^dex^).

### Dextran digestion assay

Dextran 500 (Biosynth, range of 450-550 kDa, average 500 kDa) was dissolved at 5 mg/ml in 10 mM HEPES 100 mM NaCl pH 7.5 with 66.6 µM of either wild-type or GH^dex^ E360A (single catalytic mutant) that had been purified from BL21 (DE3) *E. coli.* Each sample was incubated at 37°C / 400 rpm in an Eppendorf Thermomixer and aliquots from each reaction were taken at various time points to assess the progress of the reactions. Aliquots were immediately boiled in a heat block for 15 minutes to inactivate the enzymes, followed by a 5-minute centrifugation step at ∼17,000 RCF. Aliquots were subsequently frozen at −20°C for later thin-layer chromatography analysis. The time points were at 0 (*i.e.* before GH^dex^ was added), 1, 5, 10, 30, 60, 120, 240 minutes, and a final overnight time point (21.5 hours after initiation).

### Thin-layer chromatography

Foil-backed silica TLC sheets (Silica gel 60 F254) were cut to size and a line was drawn 1 cm from the bottom edge in pencil. Samples were loaded 1 cm apart from each other along the line by pipetting 2 µl at a time followed by drying with a hairdryer. For oligosaccharide samples, 6 µl was loaded per spot. The first and last lanes were reserved for a size marker consisting of 3 µl of 1 mM glucose, maltose and isomaltotriose, corresponding to dextran digestion products with a DP of 1, 2 and 3 respectively. The loaded sheets were then placed in a grease-sealed glass tank with ∼0.5 cm of butanol, acetic acid and water mixture at a ratio of 2:1:1. After approximately 2.25 hours, the running buffer was <1 cm from the top of the sheets, and they were removed from the tank before being carefully dried with a hairdryer. To stain the migrated sugars, the sheet was briefly submerged in a developer solution (sulphuric acid, ethanol and water; 3:70:20 v/v with 1% orcinol) for a few seconds, and then carefully dried again. Stained sheets were then taken to a heated cabinet (120°C) for a few minutes until the bands developed sufficiently.

### Isothermal titration calorimetry analysis of purified dextran utilisome lipoproteins

Using isothermal titration calorimetry (ITC), titrations were performed for GH^dex^ E360A, GH^dex^ D297A/E360A, SGBP^dex^, SGBP^dex^ Δ1-147, and SusD^dex^ on a Microcal PEAQ-ITC (Malvern Panalytical). Dextran solutions were used as the injected titrant, while the proteins were the analytes inside the ITC cell. The concentration of protein was typically 25-50 µM, while dextran varied from 1.6 mM (dextran 1.5) to 10-12.5 µM (dextran 500). Dextrans 1.5 and 3.5 were sourced from Pharmacosmos (technical grade), with larger dextrans sourced from Biosynth. The dextran concentrations were optimised experimentally to obtain as many injections covering the inflection point of the ITC curve as possible. The standard procedure started with 10 minutes of equilibration at 25°C at a reference power of 10 µcal/s set at 750 rpm of stirring speed. The first injection of 0.4 µl of the titrant (ignored in the analysis) was followed by 18 injections of 2 µl. The initial delay was 60 seconds, and injections lasted 4 seconds and were spaced with 150 second pauses. As initial heats of injection were very large, the buffer-to-buffer heats were assumed to be minimal, and the “fitted offset” option was used to subtract a constant assumed control heat from each injection.

Since dextran is highly soluble in water, and the commercially available dextrans come as powders, dialysis was not performed. Dextran solutions were made up to particular molar concentrations based on their average molecular weight. However, commercial dextrans are only purified to a certain degree and there is some uncertainty in the exact polymeric contents. For example, technical grade dextran 1.5 (Pharmacosmos) has a molecular weight range of 1200-1800. Dextrans were dissolved in the same buffer as the protein (10 mM HEPES, 100 mM NaCl, pH 7.5) and briefly heated to ∼50°C until fully dissolved, and then aspirated with a pipette or vortexed as required.

Different analyses were used on the Microcal PEAQ-ITC software. Firstly, the “one set of sites” model was used with the number of binding sites (N) was fixed to 1 (Table S2), allowing the concentration of the dextran ligand to vary from the calculated concentration (e.g. 1.6 mM of dextran 1.5) to fit the data. Protein concentration was fixed at 30 µM for both GH^dex^ mutants, 25 or 50 µM for SGBP^dex^, 41.3 µM for SGBP^dex^ Δ1-147, and 50 µM for SusD^dex^. The appropriate dextran concentration was determined through trial and error and then repeated to get at least triplicate data. Secondly, for stoichiometry analysis (Table S5) the same model was used but N was allowed to vary instead of the concentration of ligand. Binding mutations were also assessed (Table S2, N value = 1). For mutation analysis, protein concentration was fixed at 36 µM for GH^dex^ ^D297A^ ^E360A^ ^W260A^, 30 µM for GH^dex^ ^D297A^ ^E360A^ (“WT”), 25 µM for all SGBP^dex^ constructs, and 57 µM for SusD^dex^ ^Y103A^ or 50 µM for WT, SusD^dex^ ^W284A^, and SusD^dex^ ^F296A^. The appropriate concentration of dextran and protein was determined through trial and error. To control the comparisons, the dextran ligand concentration was fixed per-comparison (*e.g*. the same for SusD^dex^ WT and its variants).

### Crystallisation of GH^dex^, SGBP^dex^, SusD^dex^

After size-exclusion chromatography, purified proteins were concentrated to 10-50 mg/ml using 15 ml Amicon centrifugal filters (MWCO = 30 kDa) (Merck Millipore). Proteins were screened by sitting drop vapour diffusion crystallisation using the PACT, Structure, Morpheus I and JCSG+ screens (Molecular Dimensions), the Index screen (Hampton Research), and if the target crystallised in ammonium sulphate conditions, the AmSO4 screen was also used for hit optimisation (QIAGEN). For each of the 96 conditions of the screens, the Mosquito (SPT Labtech) deposited mixtures of protein and crystallisation solutions; 100:100 nl (1:1) and 200:100 nl (2:1) respectively. MRC 96-Well 2-Drop Crystallization plates (Molecular Dimensions) were used and sealed with ClearVue sheets (Molecular Dimensions). To optimise confirmed protein crystals, the Dragonfly (SPT Labtech) was used to mix a series of conditions based on the contents of the original crystallisation condition, typically varying the amount of precipitant and/or salt by 20% above and below the original condition. Crystallisation plates were kept at either 20 or 4°C and inspected regularly with an optical stereomicroscope for crystal formation. A polarising lens was sometimes used to check for birefringence in protein crystals.

All crystals were grown at 20°C. Apo GH^dex^ E360A was crystallised in 2 M ammonium sulphate, 0.2 M potassium sodium tartrate, 0.1 M sodium citrate, pH 5.6. GH^dex^ E360A co-crystallised with dextran 1.5 (5 mM) was grown in 1.6 M ammonium sulphate, 0.5 M lithium chloride. Wild-type GH^dex^ was crystallised in a MES-containing (0.1 M, pH 6.0) ammonium sulphate condition (1.6-2.4M). The final apo SusD^dex^ crystal was grown in 0.8M ammonium sulphate, 0.1 M citric acid, pH 5. An initial crystal was also grown in 50% w/v PEG 400, 0.2 M lithium sulphate, 0.1 M sodium acetate pH 4.5, yielding a marginally lower resolution structure. The dextran-bound SusD^dex^ structure was obtained by soaking crystals grown in the PEG 400 condition with 50 mM dextran 1.5, which caused the crystals to crack into pieces. Fragments were harvested and yielded the dextran-bound structure. Initial SGBP^dex^ Δ1-147 crystals were grown in 25% w/v PEG 4000, 0.2 M ammonium sulphate, 0.1 M sodium acetate, pH 4.6. Optimised SGBP^dex^ Δ1-147 crystals were grown based on Morpheus H8 (Precipitant mix 4 [MPD, PEG 1000, and PEG 3350] 35.5% v/v, 200 mM amino acid stock [0.2 M alanine, glutamate, lysine, serine and glycine], 100 mM system 2 buffer [0.1 M HEPES sodium salt, 0.1 M MOPS, pH 7.5]). Crystals were harvested with appropriately sized, 20 μM thick nylon loops (Hampton Research), and cryo-protected in a 4:1 mixture of reservoir solution and either 100% PEG 400 or MPD, 6 M NaCl, or 3.5 M (NH_4_)_2_SO_4_ matching the crystallisation condition. Crystals from the Morpheus screen did not need to be cryo-protected. Crystals were then flash frozen in liquid nitrogen before being stored in canes in a cryo dewar.

### X-ray crystallography data processing

X-ray crystallography data was processed using the CCP4cloud software suite[41]. A typical workflow consisted of the following. Spot-finding, indexing and integration was performed using *DIALS* [42]. After this, space group was estimated with *POINTLESS* [43], and scaling, merging and high-resolution cut-off determination with *AIMLESS* [44]. Datasets were truncated based on outer-shell statistics for CC_half_ and mean(I) / σ(I), where ideally the former was greater than 0.5 and the latter greater than 1.0, with greater importance given to CC_half_. Asymmetric unit content determination via Matthews coefficient was performed by the CCP4 implementation [45], and molecular replacement was carried out with *phaser* and *molrep* [46,47], followed by refinement using *refmac5* and *refmacat* [48,49]. *CCP4build* [50] was used for early model building and refinement where large modifications were required. Computer-assisted model building was performed in *Coot* [51] and cycles of *refmac5* refinement and model-building proceeded until the model could not be improved, primarily judged by R_free_ values and *MolProbity* [52] scores after refinement. Modelled waters (represented as red spheres) were found and checked with primarily with *Coot*. *CheckMyBlob* [53] was used to model unknown ligands into the difference map. A full list of data collection statistics, derived from the final *Refmac* refinements for each crystal structure can be found in Table S1.

### Tagging of the catalytically inactive dextran utilisome in *B. theta*

*B. theta* VPI 5482 strains were created using a previously described counter-selection protocol [9,10]. The pExchange-tdk (pEX) vector [29] was used to introduce mutant genetic sequences from *E. coli* S17 λ *pir* to the genome of *B. theta (tdk-)* through conjugation and homologous recombination. Cells that contained pExchange-tdk were then selected, resulting in either wild-type or mutant recombinants, which were confirmed by both colony PCR and Sanger sequencing of the amplified region of interest. As the *BT3087 D297A/E360A* region of interest was previously cloned by site-directed mutagenesis of the *BT3087* pET28b construct, a pEX-compatible insert (with SalI/XbaI restriction sites) was amplified using the construct as a template, and then ligated directly into pEX. Once the *BT3087 D297A/E360A B. theta* strain was confirmed, the genomic DNA of the previously generated [10]. *BT3089*-His_6_ *B. theta* strain was amplified as a pEX insert, and then the counter-selection protocol was used to introduce *BT3089*-His_6_ into the *BT3087 D297A/E360A* background strain [16]. Colony PCR using a His-tag specific primer, and Sanger sequencing of the target region both confirmed the presence of the His-tag.

### Expression and purification of the inactive dextran utilisome

Expression and purification of the inactive dextran utilisome was carried out mostly as previously described [10]. Expression of the dextran system was induced by 0.5% (w/v) Dextran 3 MW 2500 – 4000 (Biosynth), as opposed to the previously used [10] Dextran T3.5 MW 3000-4000 (Pharmacosmos). For the cryo-EM sample, 4 L of *BT3087 D297A/E360A, BT3089-His6* cell membranes were extracted using 2% DDM, followed by exchange to LMNG during IMAC. The column was first washed with 25 ml of TBS (0.15% DDM, 30 mM imidazole, followed by 25 ml of TBS (0.005% LMNG, 30 mM imidazole). For elution, 5 ml of TBS (0.002% LMNG, 200 mM imidazole) was used. After concentration with an Amicon Ultra centrifugal filter (MWCO = 100 kDa), size exclusion chromatography (SEC) was performed with a Superose 6 Increase 10/300 GL column, as the use of a Superdex 200 column is precluded with dextran-binding proteins due to strong interactions. The column was run with size exclusion buffer without detergent (10 mM HEPES, 100 mM NaCl, pH 7.5) to minimise excess LMNG in the final sample.

### Cryo-EM sample preparation and data collection

For better particle distribution and number on cryo-EM holey grids, UltrAuFoil R 2/2 grids on 200 gold mesh were chemically modified with a self-assembled layer of polyethylene glycol (PEG)[30]. The grids were first glow-discharged at 20 mA for 90 sec on each side (total of 180 sec), transferred into an anaerobic glovebox (< 2 ppm O_2_) and submerged in ethanol containing 5 mM hexa(ethylene glycol)mono-11-mercaptoundecyl ether (675105, Sigma Aldrich) for ∼24 hours. Just prior to use, the grids were removed from the glovebox and washed successively in three tubes of ethanol and left to air-dry. For vitrification, 0.5 mM of dextran T1.5 (Pharmacosmos) and dextran 3 (Biosynth) were added to 7.5 mg/ml of inactive dextran utilisome, and equilibrated for at least 30 minutes before vitrification at room temperature. Fluorinated octyl maltoside (O310F, Anatrace) was also added at its CMC (0.7 mM). 3 µl of utilisome was applied to the grid held in a Thermo Scientific Vitrobot IV plunge freezer. The chamber was kept at 4°C with 100% relative humidity. After a 3 second blot delay, a blot force of −5 was applied for 6 seconds. For data collection, a remote collection was performed on the Titan Krios II at eBIC (Diamond Light Source). 12,408 movies were collected at a magnification of 105,000x using a Gatan K3 detector, resulting in a physical pixel size of 0.831 Å. Electron dosage was 23.1 e/Å^2^, and nominal defocus values were increments of 0.2 µm between −1.0 and −2.0 μm. See Table S3 for details.

### Cryo-EM data processing of the inactive dextran utilisome

*CryoSPARC* [54] was used for processing cryo-EM movies (Table S3). Two processing pipelines were performed. For pipeline 1 (Figure S6), patch motion correction and patch CTF were first performed on imported movies, followed by “Curate exposures” to remove low-quality images. Template picker (using 2D references from earlier datasets) was used for particle picking. Most particles were removed through cycles of 2D classification and picking of the best-looking classes. Ab-initio reconstruction was performed with two output classes to separate “junk” particles from “real” ones. Iterative rounds of non-uniform (NU) refinement and 2D classification were then used to remove more junk particles while increasing the particle box size up from 128 pixels, until the refinements were not Nyquist limited, based on interpretation of Fourier-shell correlation (FSC) curves. This generated a consensus reconstruction of 273,062 particles. 3D classification was then performed on this subset and found distinct classes for “closed-closed” (CC) and “open-closed” (OC) utilisomes. CC and OC particle subsets were then non-uniform refined separately (175,481 and 97,400 particles respectively). Another round of 3D classification was used for both particle stacks. This did not improve the resolution for the CC map, but the two best classes with a fully open SusD were chosen from the 3D classification for the final OC reconstruction of pipeline 1, which was used for model building. Using the consensus particle stack, 3D variability analysis [31] was performed which led to the SusD^dex^ hinging movie (see Figure S8, Movie S1 and Table S6).

For pipeline 2 (Figure S7), the dataset was reprocessed for compatibility with the Cryosparc “reference-based motion correction” job, which can lead to improvement in map quality. Particle numbers in Table S3 refer to maps output from pipeline 2. Patch motion correction and patch CTF were applied to the movies, followed by “Curate exposures” to remove low-quality exposures. Particles were blob picked (140-160 Å), extracted into 400 pixel boxes, and then subjected to several rounds of 2D classification. Ab-initio reconstruction and homogenous refinement (1 class) were performed on the remaining 481,934 particles, followed by ab-initio (5 classes) and homogenous refinement, followed by ab-initio and heterogenous refinement (5 classes each). This yielded two noise classes, classes for OC and CC utilisomes, and one class of the core SusCD complex. The OC (∼150,000 particles) and CC (∼215,000 particles) classes were independently non-uniform (NU) refined to 3 and 2.9 Å, respectively. Reference-based motion correction and global and local CTF refinement was performed independently on OC and CC particle stacks, resulting in nominal resolutions of 2.5 and 2.3 Å for each state, respectively. To reduce the compositional and conformational heterogeneity of these consensus reconstructions, 3D classification (10 classes) was then performed with “force hard classification” on. For the CC stack, the best class with density for all eight protein units was used for final model building, with 116,104 particles producing a 2.5 Å reconstruction. The OC stack contained more conformational heterogeneity with classes containing different degrees of open SusD^dex^. Classes 1, 5 and 9 had a fully open SusD^dex^ and refined to 2.7 Å when their particles were combined, but the density for one copy of GH^dex^ was poor. The best octameric fully open class (class 1, 21,574 particles) was refined to 2.9 Å and used for final OC model building.

Unsharpened maps, mirrored into the correct hand, were used for model building. The cryo-EM structure of SusC^dex^ (PDB ID: 8AA4) and X-ray structures of GH^dex^ and SusD^dex^ reported in this study (PDB ID: 9SJG, 9SJK) were rigid body docked into the density and refined with several rounds of *Phenix* [55] real space refinement and semi-automated model building in *Coot* [51]. An Alphafold2 model of SGBP^dex^ was also docked, but only the NTD was sufficiently resolved in the map, so the CBM domains were deleted from the model. Dextran ligands were built using glucose (GLC) monomers and O6 to C1 were covalently linked with the *Coot AceDRG* plugin [56]. *Privateer* [57] was used to provide glycan-specific restraints for further refinement. Finally, global and local *Isolde* [58] simulations were used to validate the model composition and lessen clashes and geometric outliers, and its output was supplied to *Phenix* for a final refinement.

### Accession numbers

The OC and CC cryo-EM structures (Table S3) were deposited under entries 9SJE and 9SM2 (PDB) and EMD-54945 and EMD-55024 (EMDB), respectively. Crystal structures (Table S1) were deposited in the PDB under accession codes 9SJG (GH^dex^ WT), 9SJH (GH^dex^ E360A, apo), 9SJF (GH^dex^ E360A, with dextran), 9SJI (SGBP^dex^ Δ1-147), 9SJK (SusD^dex^, apo), and 9SJJ (SusD^dex^, with dextran).

## Acknowledgements

The research of BvdB was supported by a Wellcome Trust Investigator award (214222/Z/18/Z), with funding for MFs PhD studies provided by Newcastle University. We thank the Newcastle University EM Research Services for assistance with the generation of negative stain EM images. We also acknowledge York Structural Biology Laboratory (YSBL) for assistance with cryo-EM screening and data collection, Prof. Jamie Blaza for assistance with cryo-EM sample preparation, and Dr. Alan Cartmell for assistance with glycan model building. We would also like to thank Diamond Light Source for access and support of the cryo-EM facilities (proposal bi28576-19) at the UK national electron Bio-Imaging Centre (eBIC), as well as for synchrotron beamtime (proposal mx24948) with assistance from the staff of beamlines I03, I04 and I24 with crystal testing and data collection.

## Author contributions

MF – performed cloning, protein purification, ITC, X-ray crystallography and cryo-EM, manuscript drafting and editing.

AS – assistance with cryo-EM data collection and molecular modelling.

AB – X-ray synchrotron data collection, assistance with cryo-EM data collection and molecular modelling, management of Newcastle Structural Biology Facility.

BvdB – supervision of MF, funding acquisition, project conceptualisation, assistance with cryo-EM and crystallography sample preparation, manuscript drafting and editing.

