## Supplementary data for "Structural and functional characterisation of the dextran utilisome from *Bacteroides thetaiotaomicron*"

**Supplementary Information**

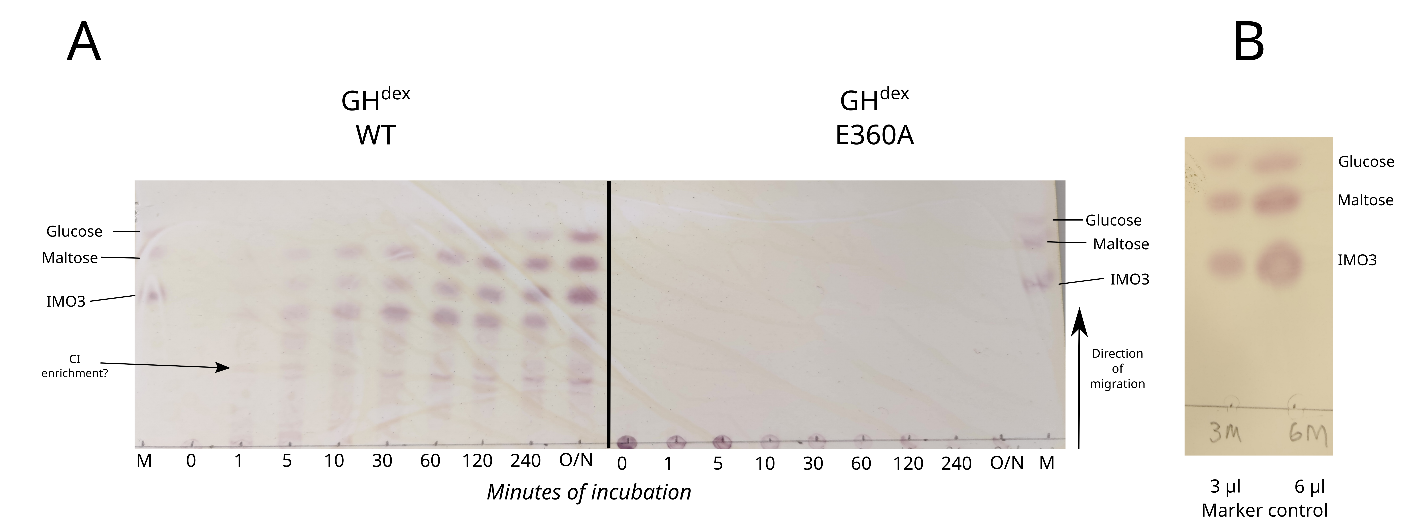

**Figure S1. Assessing the dextran degradation activity of GH^dex^ wild-type and E360A. A.** thin layer chromatography plate showing glycan reaction products from WT GH^dex^ (left) or GH^dex^ E360A (right) digestion of dextran 500. Stained saccharides are separated by size, where smaller saccharides migrate the furthest upwards. On the x-axis are time points of the dextran digestion assay and an overnight sample taken after ~19 hours (O/N). The loaded marker (M) was 3 µl of a mixture of glucose, maltose and isomaltotriose (IMO3) at 1 mM each. (B) Marker control to check for appropriate amount of sugar to load and to confirm the three bands that were not as clearly seen on the main TLC plate. An enrichment of a larger glycan was noted over the time course, which may result from the secondary activity of GH^dex^ to catalyse the formation of cycloisomaltosaccharides (CI) [1].

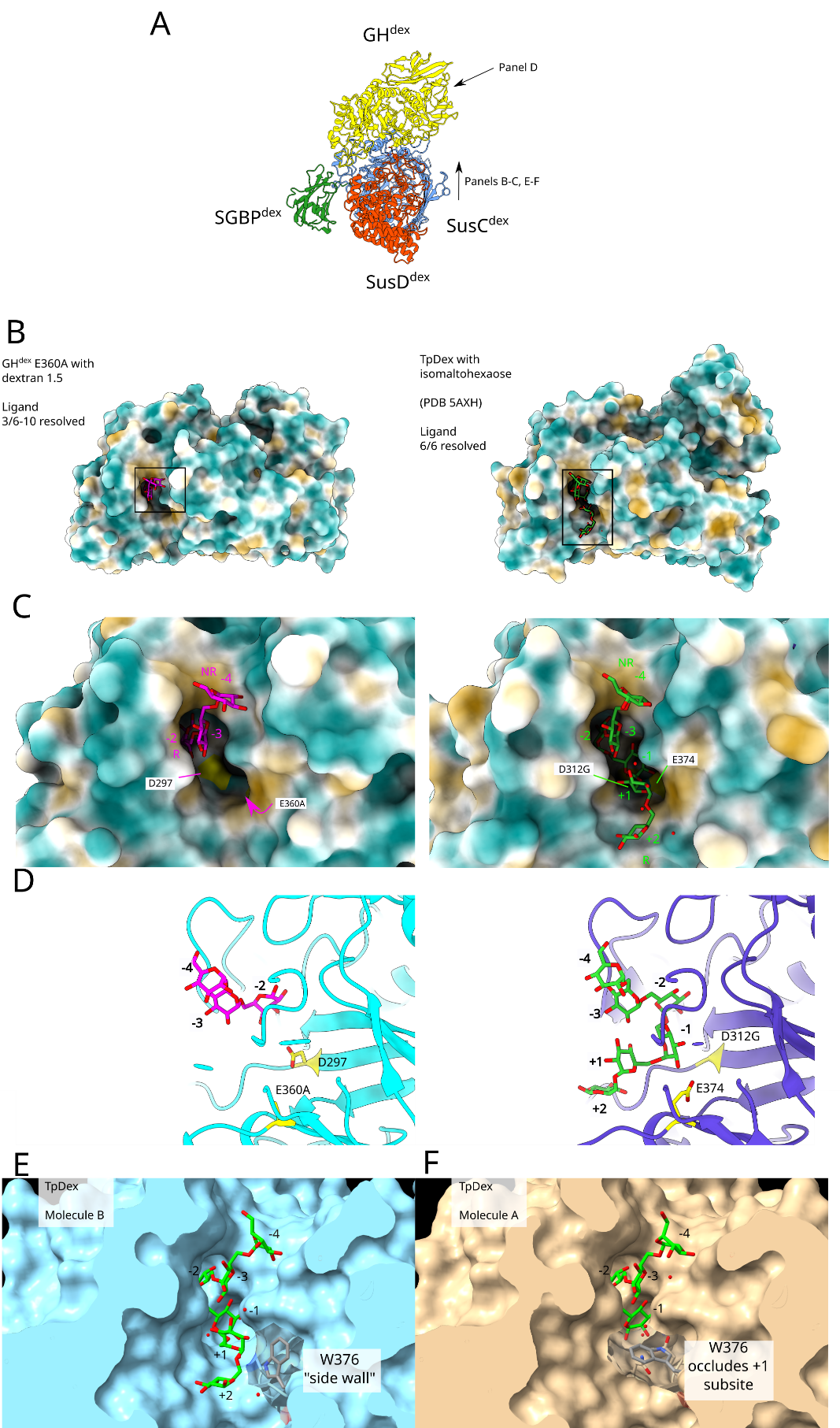

**Figure S2. Comparison of dextranase structures. A.** Orientation panel showing a top-down view of the dextran utilisome complex. **B-D.** Surface and atomic model views of the co-crystal structure of GH^dex^ E360A (left) with dextran 1.5 (DP = 6-10) and the same views of thermophilic dextranase *Tp*^dex^ D312G (right) complexed with isomaltohexaose (PDB 5AXH). B and C show the enzymatic site from the SusD^dex^ in the complex (in orange-red), while panel D slices through the GH^dex^ (in yellow) to present a side view. In B and C, the enzymes are hydrophobic surfaces (where hydrophilic regions are coloured in turquoise and hydrophobic are in tan). Resolved glycan residues (as well as how many were in the supplied glycan) are listed in B. Dextran-based ligand is shown in magenta, and the Tp^dex^ ligand is shown in lime green. Catalytic residues (or alanine or glycine point mutations) are shown in yellow and labelled. Waters are shown as red spheres. **E-F.** Crystallographic molecules B and A in the *Tp*^dex^ structure show different conformations of the side-wall Trp376 residue. In molecule B, W376 is in the side wall conformation, whereas in molecule A W376 is bent into the binding site, occluding subsite +1. This movement may reflect a mechanism where hydrolysis products are ejected from the enzyme [2].

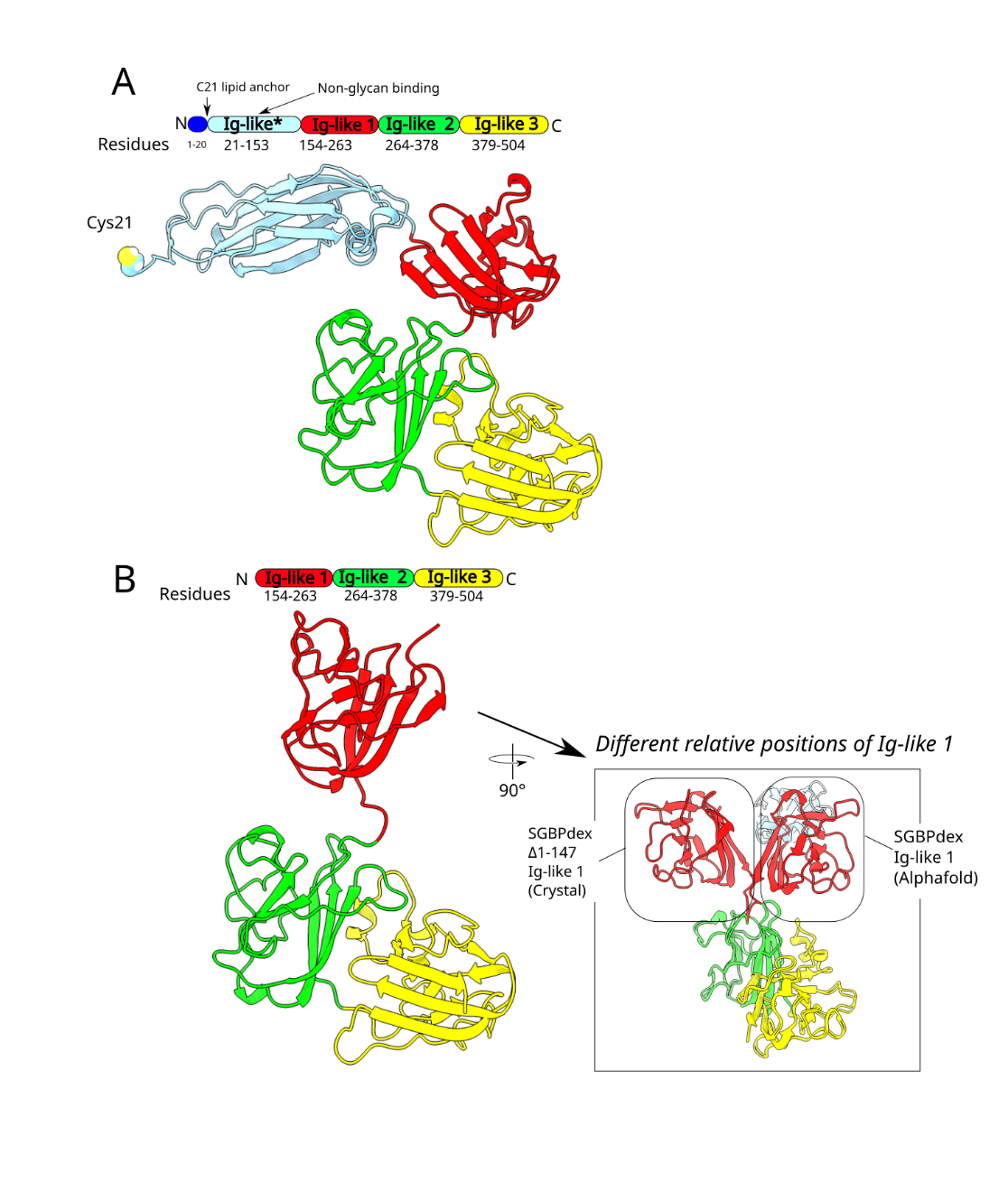

**Figure S3. Comparison of Alphafold2 models and experimental structures of SGBP^dex^. (A)** the full length AlphaFold model and **(B)** the crystal structure of the N-terminally truncated SGBP^dex^ Δ1-147. The protein models are coloured according to their domains, including an N-terminal signal sequence that is cleaved from the final protein (dark blue), the Ig-like N-terminal domain (light blue), and three carbohydrate binding modules (CBMs; red, lime green and yellow). The first structured domain in the wild type protein (Ig-like) has been removed in the Δ1-147 truncated construct, since this domain is not predicted to have a substrate binding function. The other three domains are predicted carbohydrate binding modules (CBM 1-3) based on comparison with studies on the homolog SusF [3]. In the crystal structure for SGBP^dex^ Δ1-147, the linking residues between CBM 1 and 2 angle CBM 1 towards the viewer, whereas in the Alphafold model CBM 1 is angled into the background (see boxed inset panel B for a comparison). However, when aligning the CBMs of the Alphafold and crystal structure separately, CBM 1 and combined CBM 2/3 give Cα-RMSD values of 0.8 and 0.6 Å respectively.

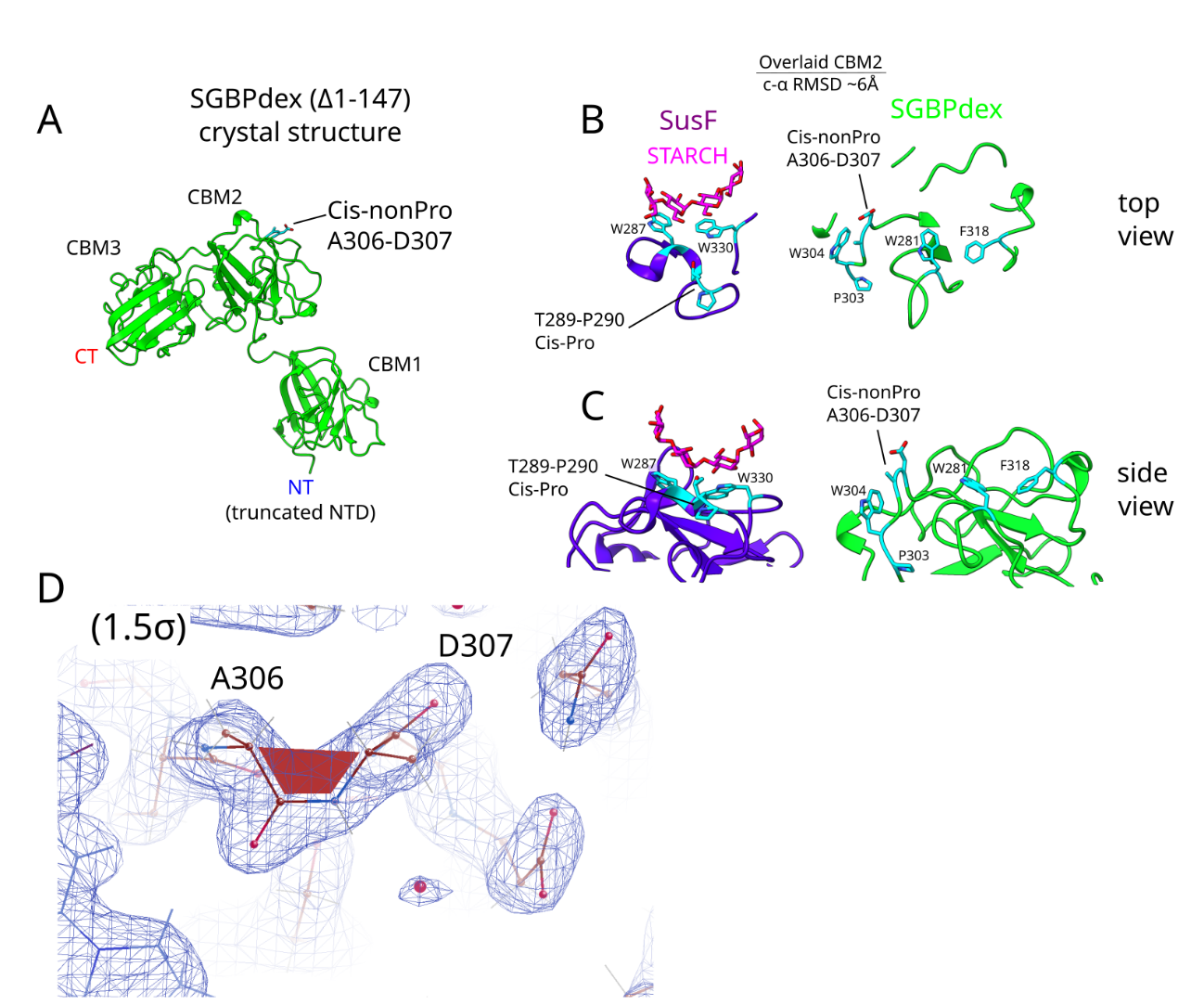

**Figure S4. A non-proline cis peptide on the surface of SGBP^dex^ CBM 2 may contribute to its putative dextran binding site.** **A.** The crystal structure of SGBP^dex^ (Δ1-147) with the labelled cis-nonPro on the surface of carbohydrate binding module 2 (CBM2). **B-C**. SusF (PDB 4FE9) and SGBP^dex^ crystal structures were overlaid and then tiled to show similar regions side by side; top views are in panel B and side views in panel C. The starch ligand that binds to SusF CBM2 interacts with two aromatic surface residues (W287, W330), which resemble the putative dextran binding site on CBM2 (W281, F318, W304). The nearby D307 involved in the cis-nonPro peptide bond may help stabilise bound dextran through hydrogen bonding. **D.** Electron density of the A306-D307 cis-nonPro peptide bond (2Fo-Fc map at 1.5 σ).

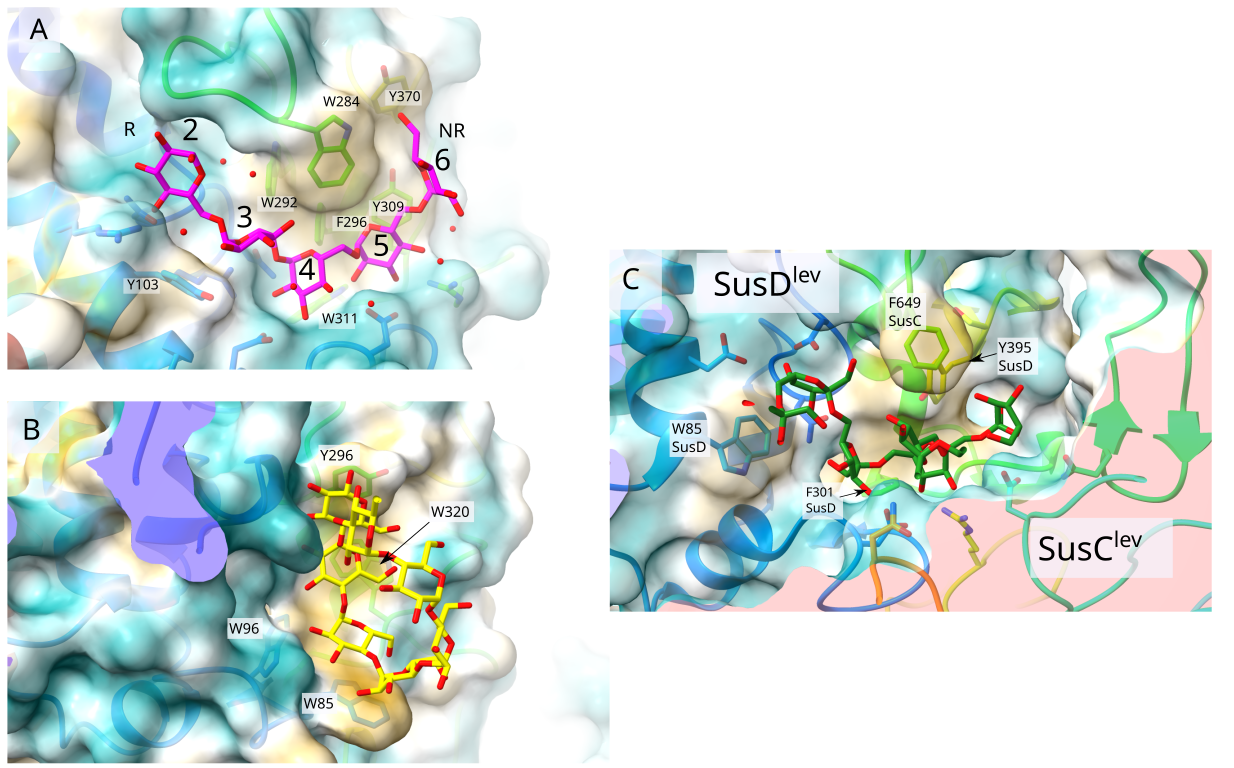

**Figure S5. Comparison of SusD binding sites.** Substrate-bound crystal structures for SusD^starch^ (**B**, PDB 3CK9) and SusCD^lev^ (**C**, PDB 6Z9A) were overlaid onto the SusD^dex^ crystal structure (**A**) using ChimeraX matchmaker. The position of the central aromatic residue SusD^dex^ Trp284 resembles SusC^lev^ Phe649, which is provided by the SusC^lev^ loop 7. The proteins are viewed towards their ligand binding regions (Figure 3 B-C, main text). SusD^dex^ is shown as a translucent surface coloured by hydrophobicity where turquoise is hydrophilic, white is neutral and tan is hydrophobic. The IMO5 that was resolved sufficiently for modelling is shown as sticks coloured in magenta with red representing oxygens. Glucose residues of SusD^dex^ substrate are labelled according to subsite numbering determined by cryo-EM (Figure 5, main text). Starch and levan oligosaccharides are coloured yellow and dark green respectively. The bound ligand in SusD^starch^ is maltoheptaose (IMO7). Ordered waters are displayed as red spheres. Aromatic residues near the ligands are labelled.

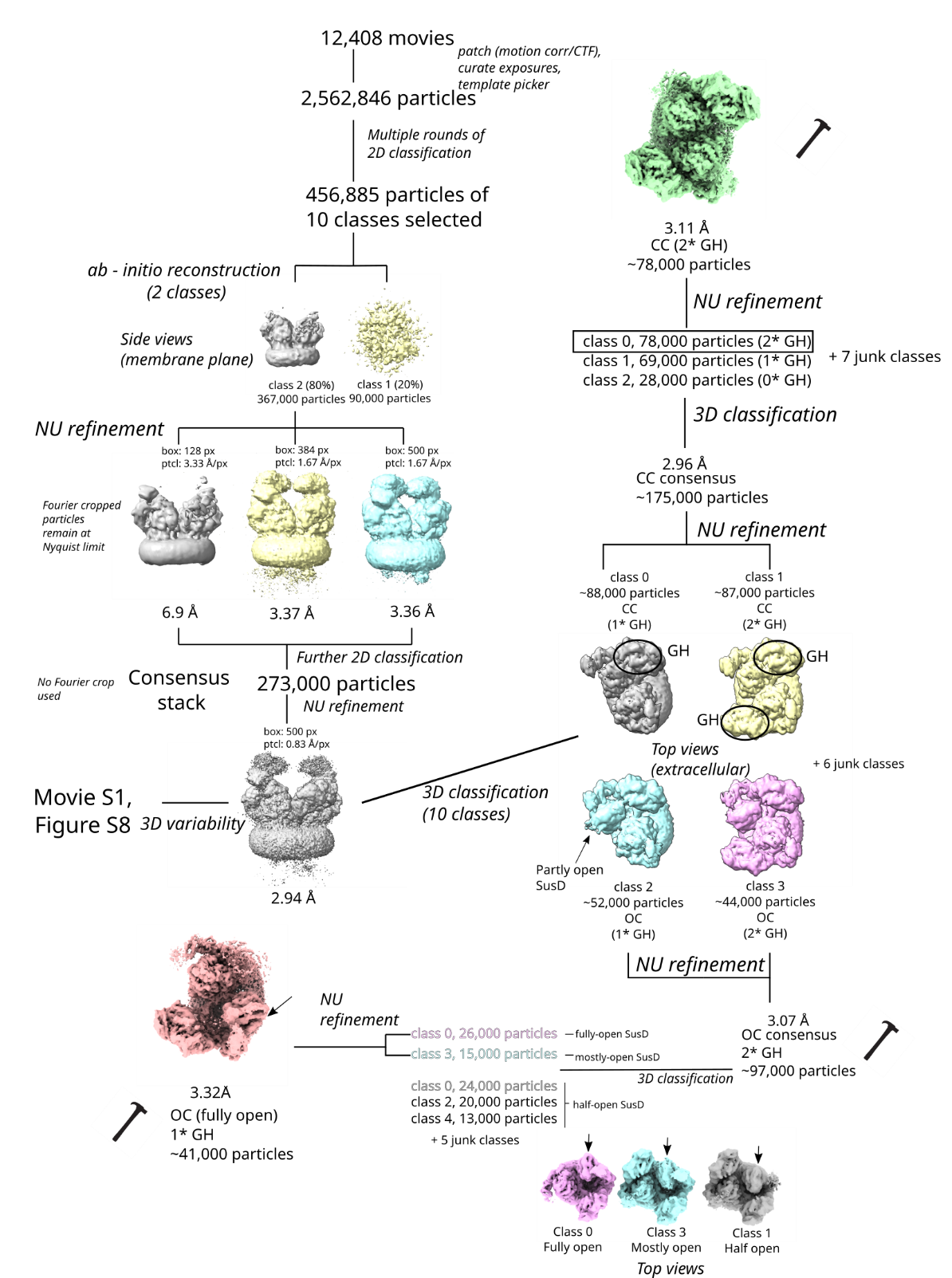

**Figure S6. Processing diagram (pipeline 1) for the closed-closed (CC) and open-closed (OC) dextran utilisome.** Maps marked with hammers were used for early-stage model building. Arrows pointing to maps indicate the SusD^dex^ in motion. Circled maps indicate the presence of a GH^dex^ subunit. Final maps were produced through a separate processing of the same dataset (Figure S7; pipeline 2). The consensus reconstruction of pipeline 1 (~273,000 particles) was used for 3D variability which generated Movie S1 and the frames in Figure S8.

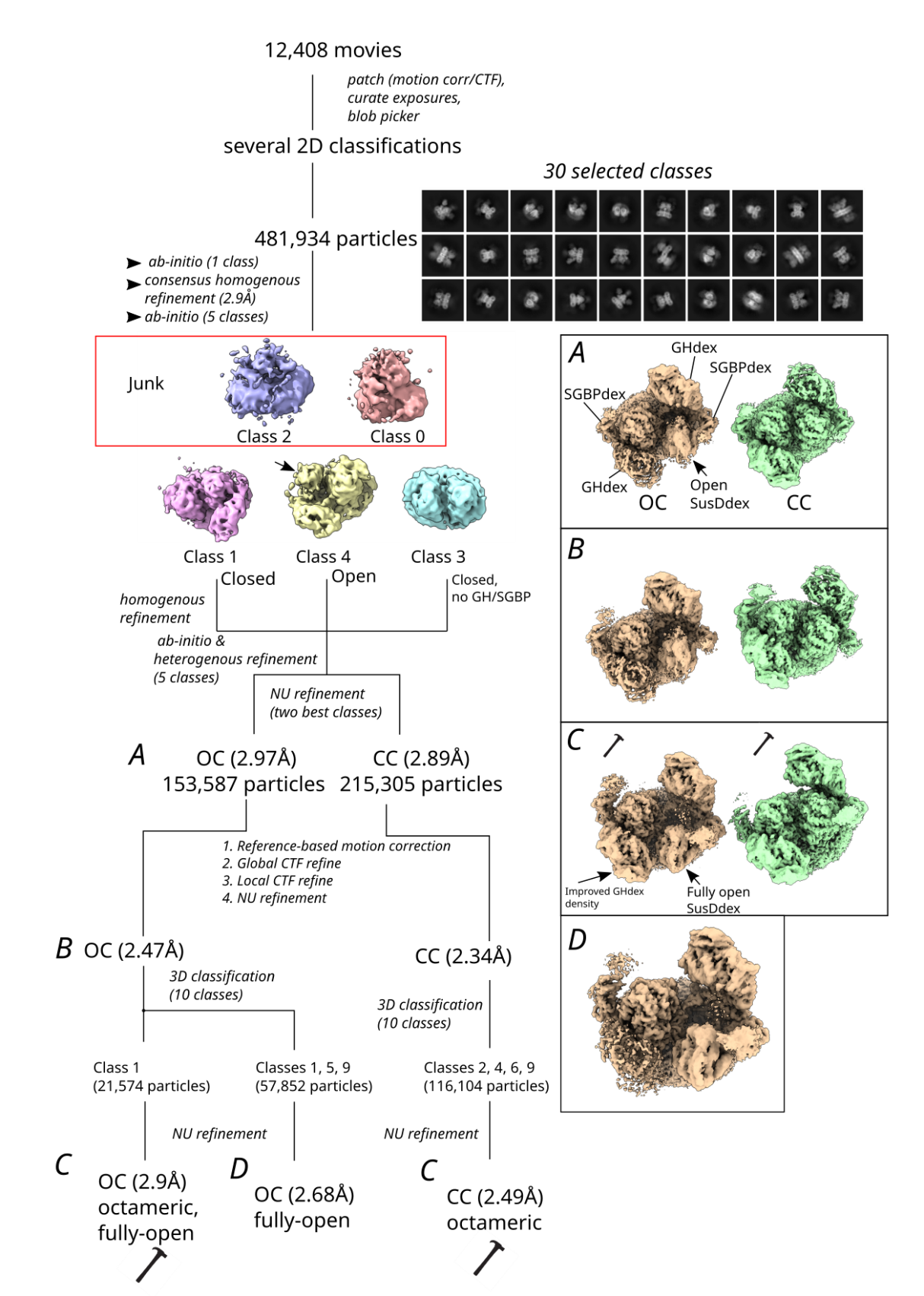

**Figure S7. Processing diagram (pipeline 2) for generation of the final closed-closed (CC) and open-closed (OC) dextran utilisome maps.** The arrow on ab-initio class 4 indicates the open SusD^dex^ lid. Top views of the maps at different processing stages **(A-D)** are shown on the right in boxes; OC is in tan and CC is in green. Maps in panel C (marked with hammers) were used for model building. Map D was generated using multiple classes that appeared to be fully open, but due to poorer GH^dex^ density this was not used.

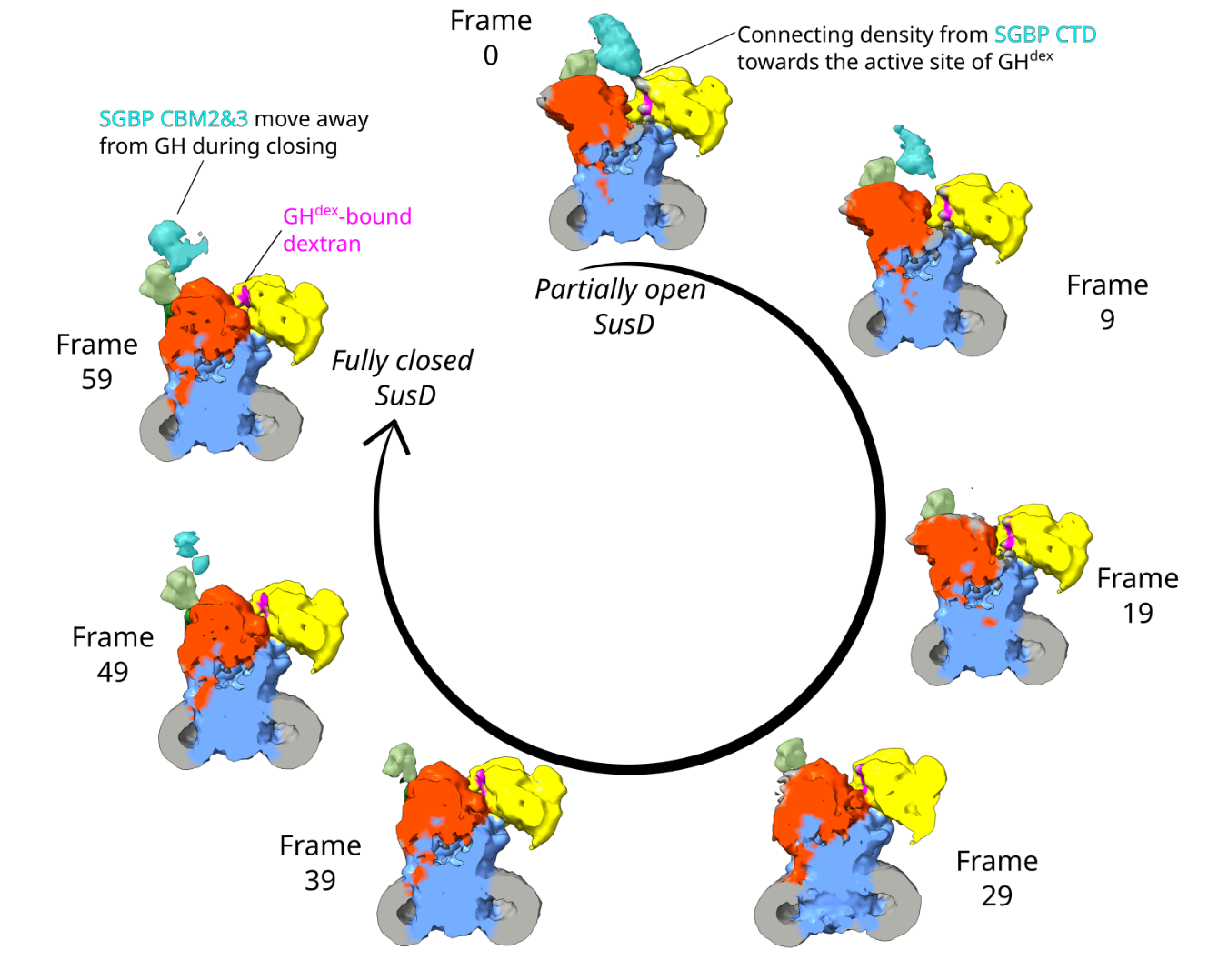

**Figure S8. Conformational cycling in the dextran utilisome.** Seven frames of the dextran utilisome transition from an open SusD^dex^ state (frame 0) to the closed state (frame 59) are shown, solved using CryoSPARC 3D variability analysis [4]. The maps have been slabbed so that one copy of the protein tetramer (SusCD^dex^, GH^dex^, SGBP^dex^) is shown from the direction of the hidden second copy. To colour the map by subunit, the CC dextran utilisome model was docked into the map of each frame as two models. The OC model was not used for docking into the open frames (e.g. frame 0), as it had SusD^dex^ forming a larger opening than the states solved by 3D variability. The first model included GH^dex^, SusC^dex^ and the SGBP^dex^ NTD, while the second model docked just SusD^dex^ to account for its variable positioning between different frames. To aid in the visualisation of the unmodelled SGBP^dex^ CBMs, these were coloured by image processing software; CBM1 in light green and CBM2/3 in cyan. IMO8 and IMO4 in the SusCD^dex^ cavity were deleted and only the GH^dex^ ligand was kept due to its relatively static position throughout the motion. Density attributed to CBM2/3 (C-terminal domain) is observed transiently between frames 0-9 and 49-59. In frames 0-9, these domains appear to contact the GH^dex^ binding site, while in frames 49-59, CBM2 and 3 was resolved on top of the closed SusD^dex^. This figure is also shown as a 20-frame transition in Movie S1.

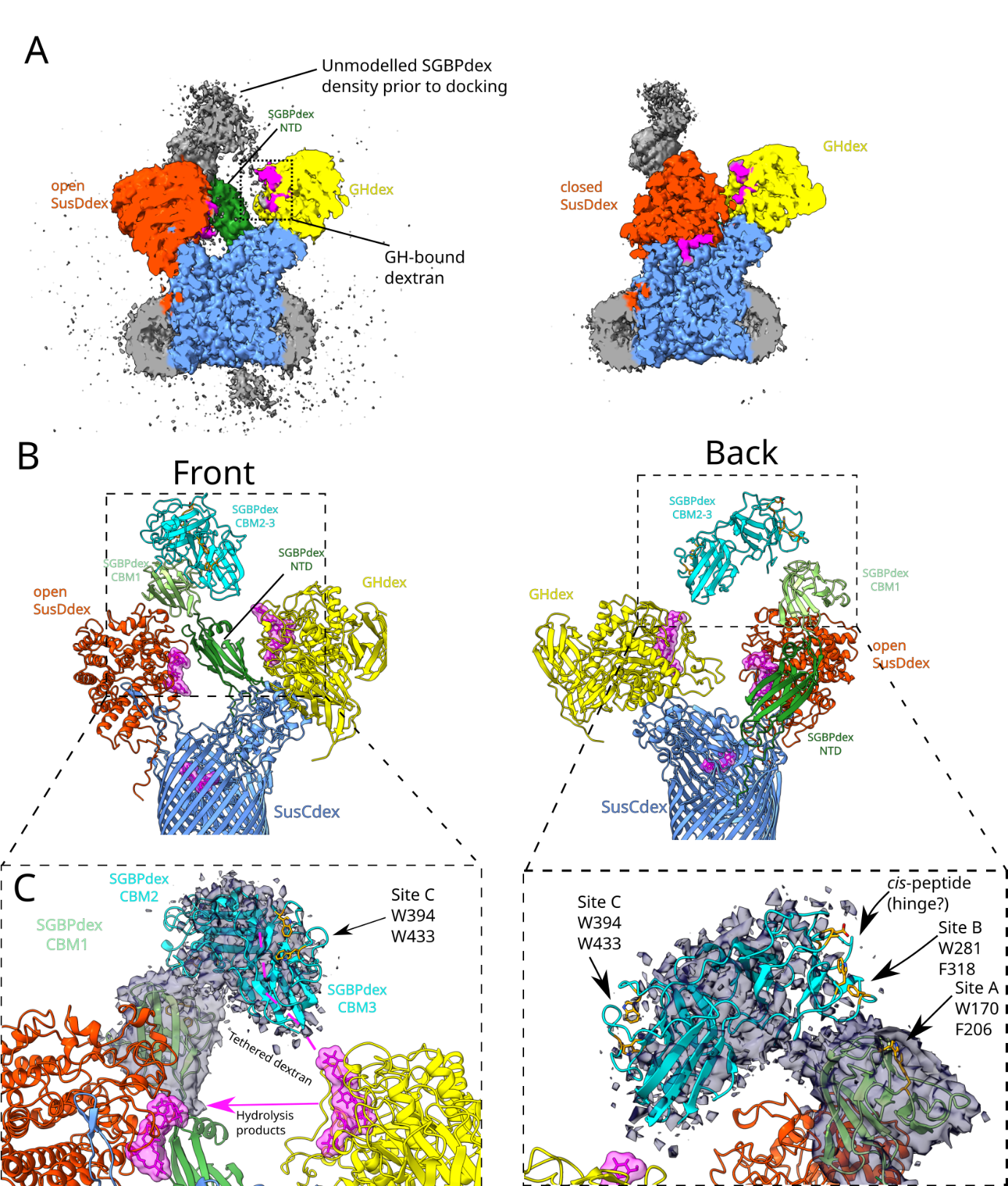

**Figure S9. SGBP^dex^ density in the OC and CC dextran utilisome maps. (A)** Side overview comparison of the open (left) and closed (right) states. The maps are viewed from inside the utilisome (i.e. from the symmetry related copies of the complex) and are both contoured at 2.5 standard deviations from the mean. Although not shown in this Figure, when taking both copies of SGBP^dex^ into account, more density of the subunit is visible in the OC map. SusC^dex^ is blue, SusD^dex^ is orange-red, SGBP^dex^ NTD (1-152) is forest green, GH^dex^ is yellow and its dextran substrate is magenta. **(B)** Front and back views of SGBP^dex^ CBM1 (light green) and SGBP^dex^ CBM2/3 (cyan) docked into the OC map. For simplicity, the closed copy of SusC^dex^ and its lipoproteins are not shown. An Alphafold2 [5] model of full-length SGBP^dex^ was split into two domains; SGBP^dex^ 153-261 (CBM1) and SGBP^dex^ 263-504 (CBM2/3). These were fit into the map in plausible orientations with respect to the N- and C-terminus and the density of the OC map. **(C)** Close-up views of the SGBP^dex^:GH^dex^:SusD^dex^ interface (left) and the docked location of SGBP^dex^ subunits (right). The map is displayed within a 5 Å radius from docked SGBP^dex^ CBM domains. The OC SGBP^dex^ density points towards the GH^dex^ (yellow) binding site. In the right panel, pairs of surface-exposed aromatic residues corresponding to putative binding sites A, B and C are shown as sticks and coloured orange. The cis non-Pro residues Ala306-Asp307 in CBM2 are also displayed.

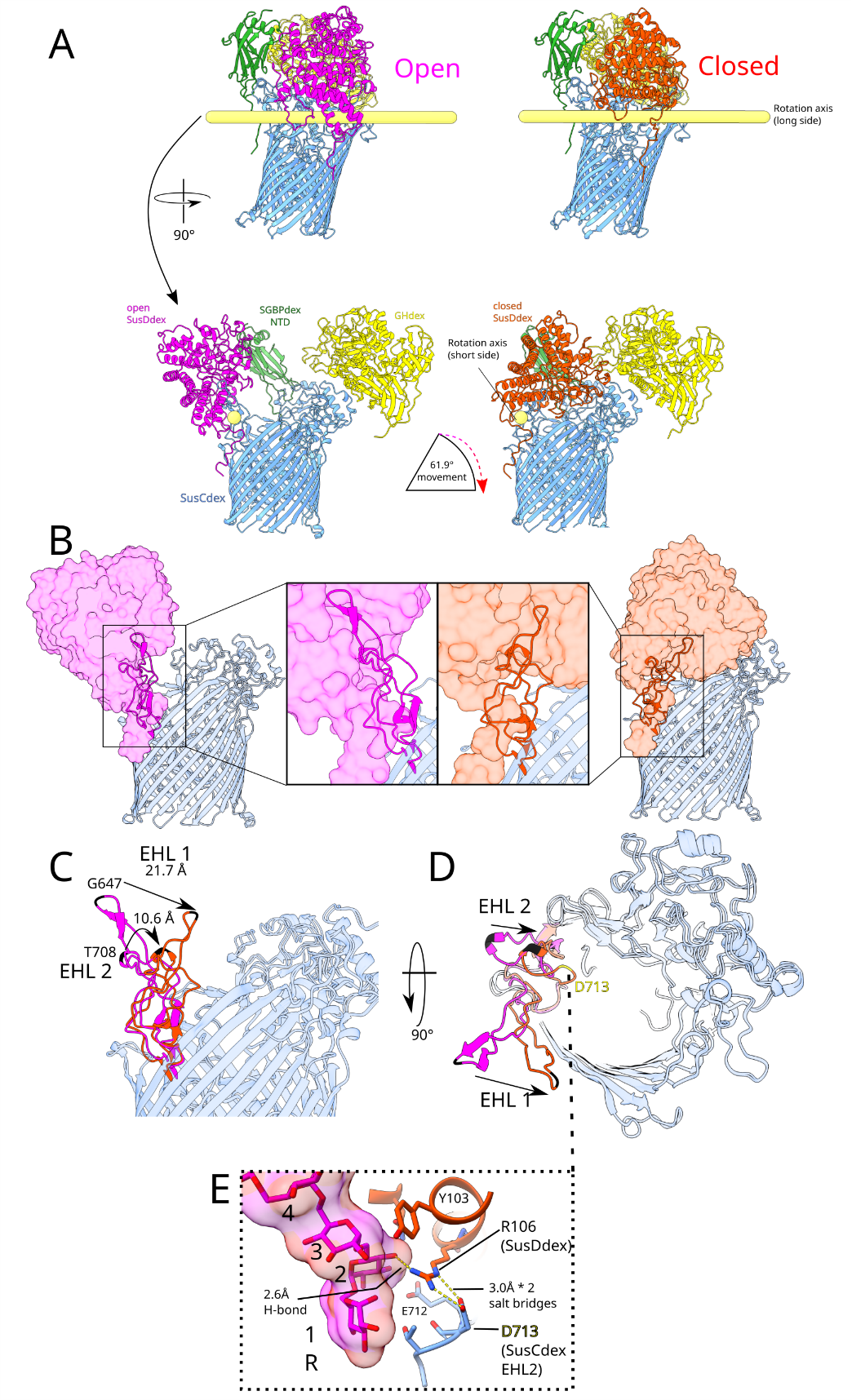

**Figure S10. Conformational changes during the transition between the open and closed dextran utilisome.** For simplicity, only one four-component complex (GH^dex^:SGBP^dex^:SusD^dex^:SusC^dex^) of the two that make up the full utilisome is shown. Dextran ligands have been hidden from view. **A.** The top panels show open and closed models where SusD^dex^ is hinging outwards towards the observer. The bottom panels show a side-rotated view along the rotation axis (yellow tube). Open SusD^dex^ is magenta, and closed SusD^dex^ is orange-red. ChimeraX “measure rotation” function was used to compare open and closed SusD^dex^. **B.** Two extracellular hinging loops (EHL) of SusC^dex^, coloured matching their SusD^dex^, contact SusD^dex^ and undergo a conformational change between open and closed utilisome states. **C-D.** Overlaid open and closed SusCD^dex^, with SusD^dex^ shown as a surface and SusC^dex^ as a cartoon. EHL1 (SusC^dex^ 627-667) and EHL2 (SusC^dex^ 697-722) are shown in full colour, while other parts of the complex are translucent. Gly647 and Thr708, at the tip of EHL1 and 2 respectively, are used to measure the movement of each loop between the open and closed states. **E.** Salt bridges (yellow dotted lines) between EHL2 residue SusC^dex^ Asp713 and SusD^dex^ Arg106, likely stabilise the CC state. Arg106 also forms a hydrogen bond with subsite 2 of the SusD^dex^ bound dextran.

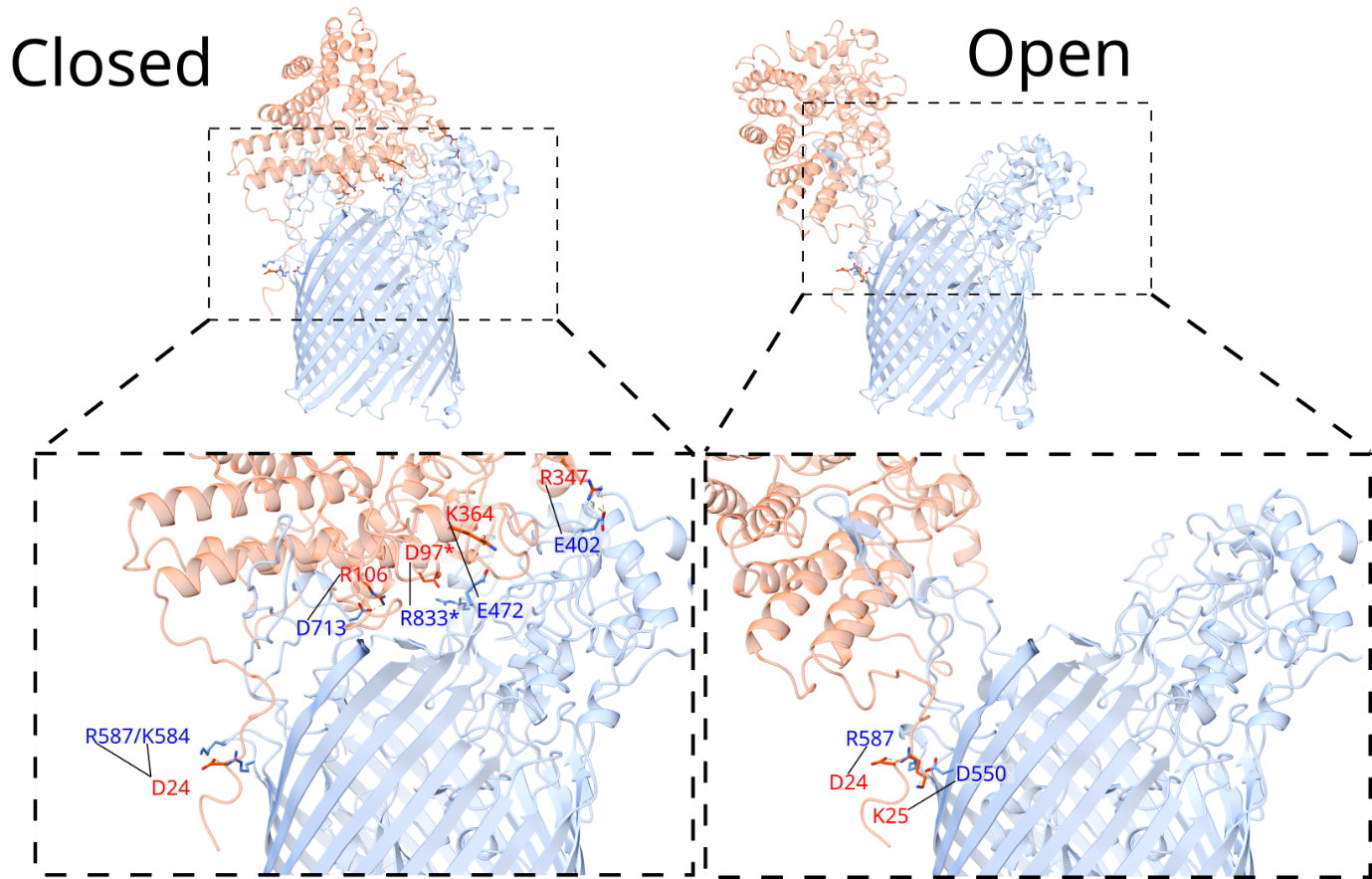

**Figure S11. SusCD^dex^ salt bridges in the closed and open states.** SusC^dex^-SusD^dex^ salt bridges were identified by the PDBePISA webserver [6] (Spreadsheet S1) and then displayed in ChimeraX with side chains shown. SusD^dex^ and SusC^dex^ residues are labelled in red and blue, respectively. The D97-R833 interaction (marked by asterisks) was included in the PISA analysis but ChimeraX did not corroborate this with the “H-bonds” function when finding salt bridges on default settings.

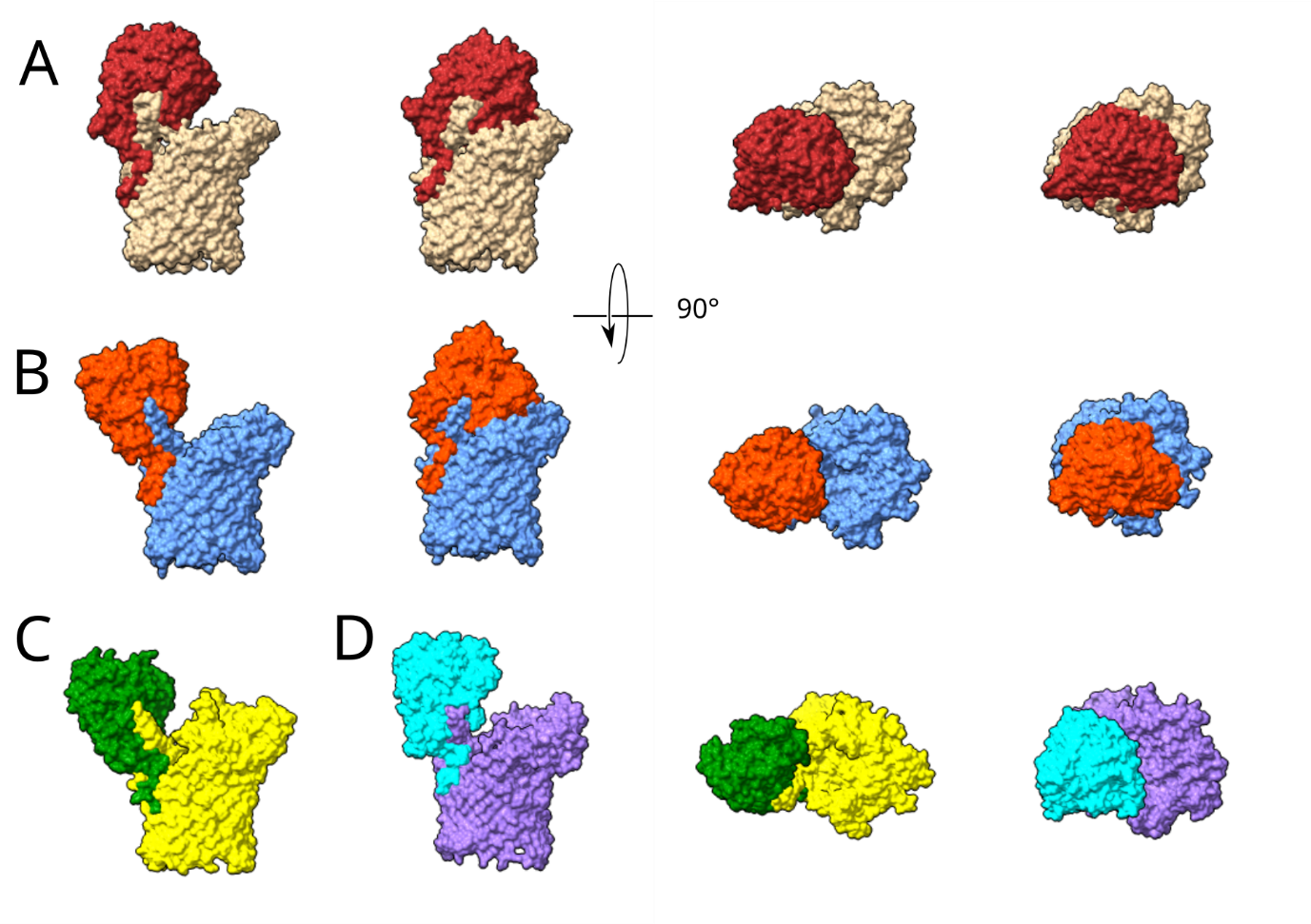

**Figure S12. Comparison of SusCD "pedal bin" apertures**. All SusC homologues were aligned on SusC^lev^ (PDB code: 6ZLT). **A.** Partly-open SusCD^lev^ (6ZLT) and closed SusCD^lev^ (6ZLU). SusD is brown while SusC is tan. **(B, this study)** Fully-open SusCD^dex^ and closed SusCD^dex^. SusD is orange red while SusC is cornflower blue. **C-D.** The open states of ButCD, a ubiquitin-homologue importer from *Bacteroides fragilis* (**C**, 8YPU) and RagAB, a peptide importer from *Porphyromonas gingivalis* (**D**, 6SML). The SusD homologues are green and cyan, while the SusC homologues are yellow and purple.

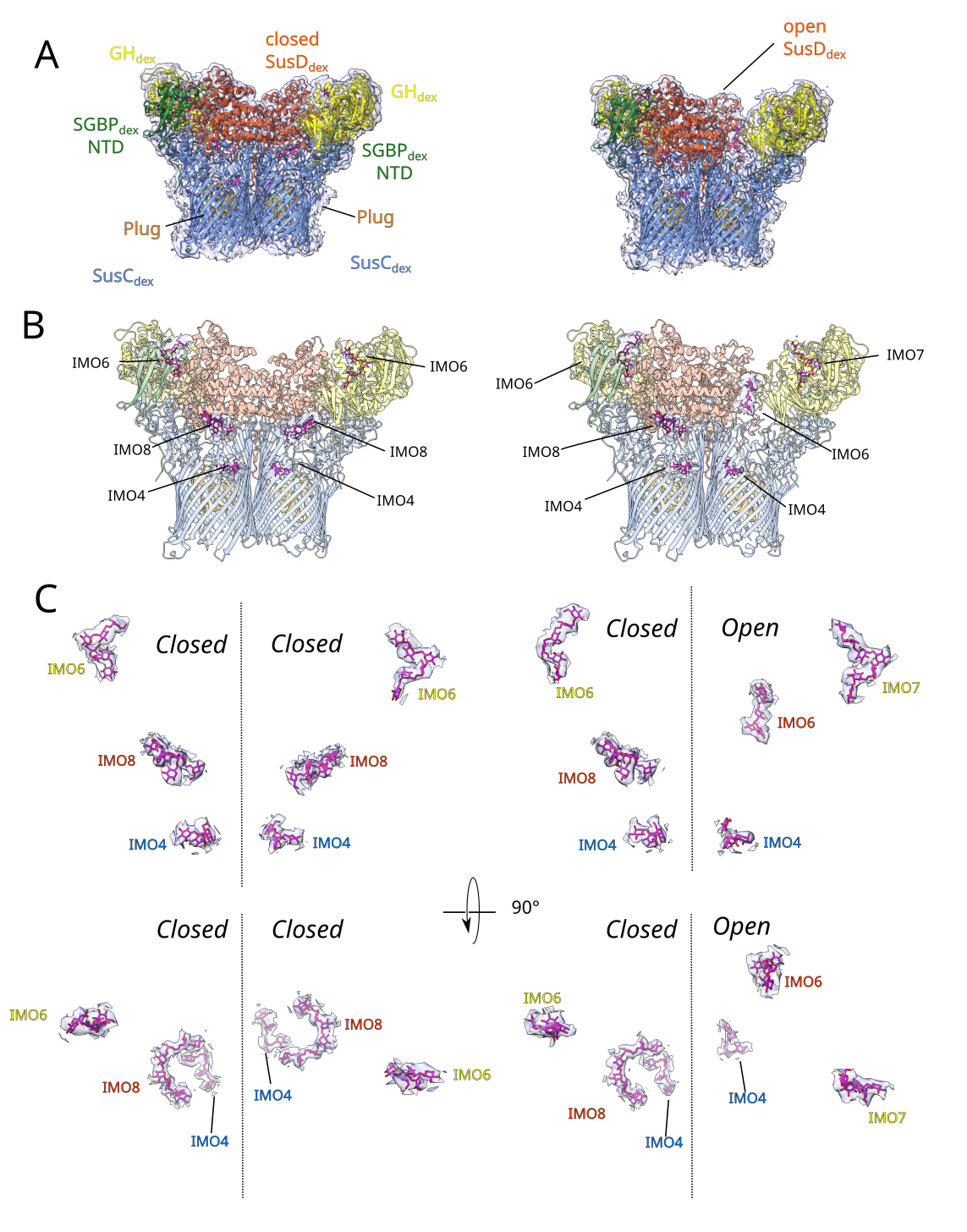

**Figure S13. Model-in-map views of the closed-closed (left) and open-closed (right) cryo-EM maps. A-B.** Models in transparent maps and the regions of the IMO ligand density. These maps were zoned to 5 Å around the whole model (A) or the ligand density (B). **C.** A magnified view of the ligands from the same view as (A/B), as well as a 90-degree rotation resulting in an extracellular view, where GH^dex^-associated IMO are in the foreground and the SusC^dex^ plug domain-associated IMO are in the background. IMOs are labelled with the colour of the subunit to which they are associated.

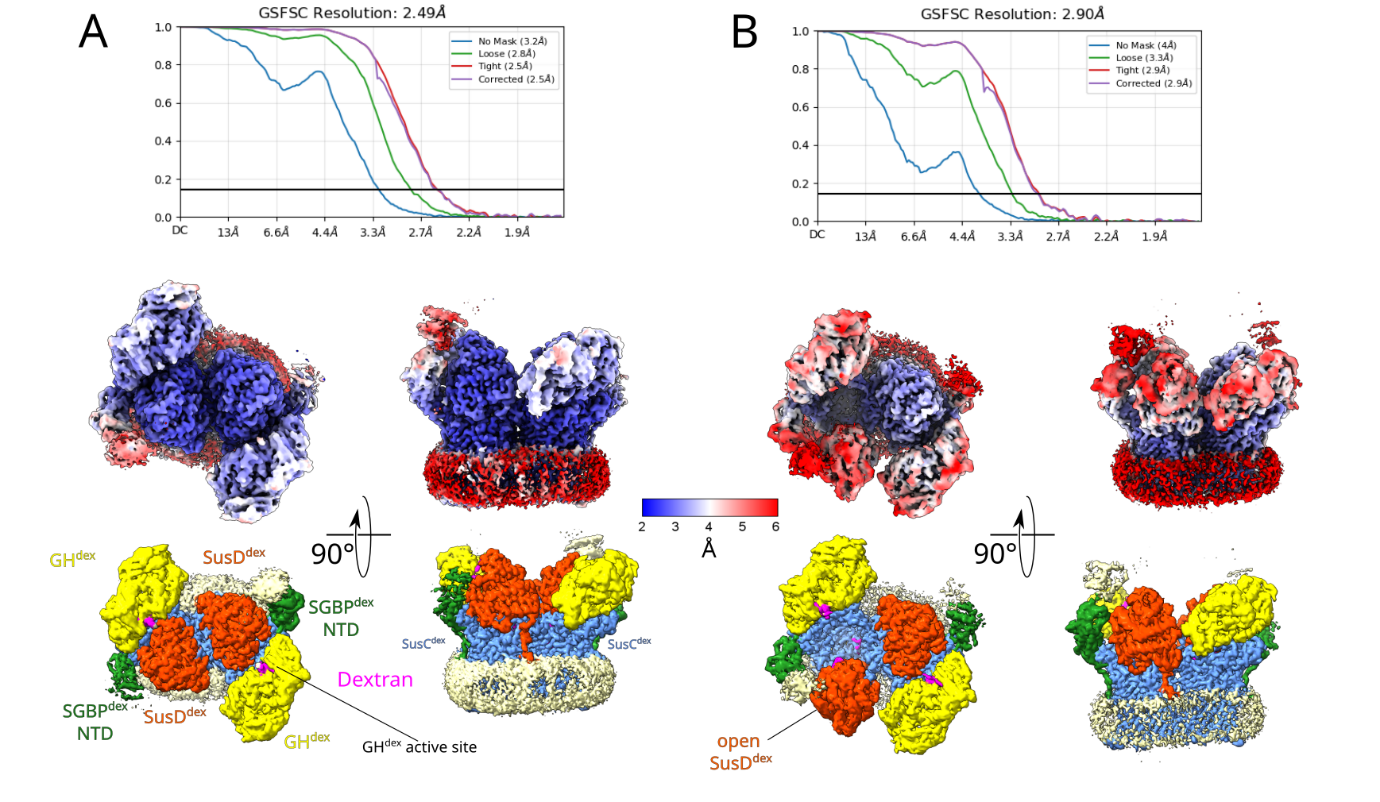

**Figure S14. The CC (A) and OC (B) dextran utilisome; Fourier-shell correlation curves, local resolution maps and maps coloured by modelled subunit.** The global resolution of the maps was estimated using the typical refinement FSC cutoff of 0.143. Local resolution was estimated in CryoSPARC, using a FSC cutoff of 0.5, and mapped from 2 to 6 Å ranging from blue to red.

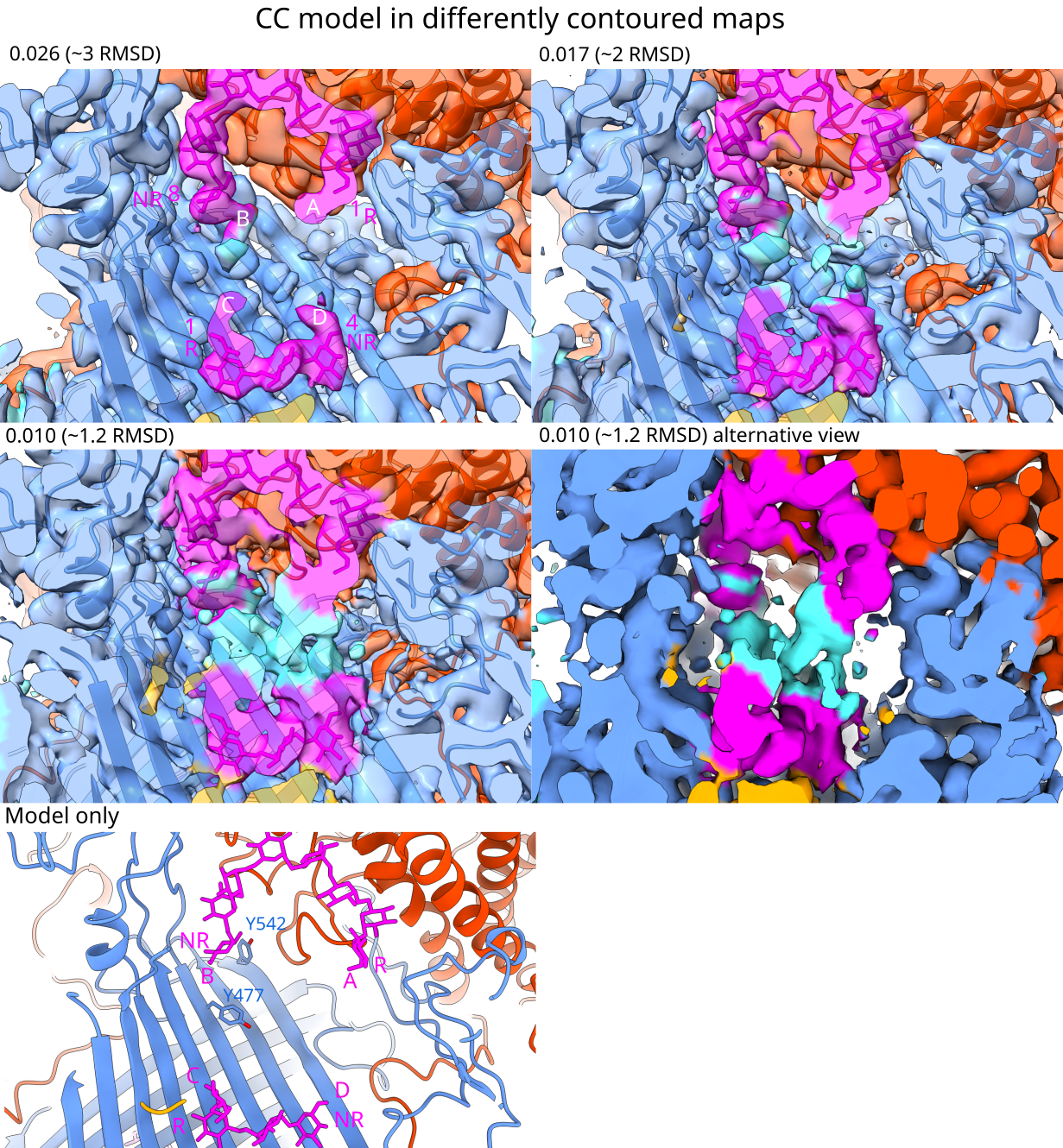

**Figure S15. The SusCD^dex^ cavity of the CC model loaded with IMO at decreasing contours.** The CC map was coloured cyan, and then coloured by protein subunit (radius around model = 5 Å, the plug domain is orange, SusC^dex^ is blue, IMO are magenta, and SusD^dex^ is orange red). The lowest contour is also shown as opaque density (without the model) for clarity. In the first panel, terminal subsites of the IMO are numbered according to the CC model (Figure 5B). In the first and last panels, the termini of the SusD^dex^–associated IMO8 (A, B) and the SusC^dex^–associated IMO4 (C, D), as well as the non-reducing (NR) and reducing (R) ends, are labelled.

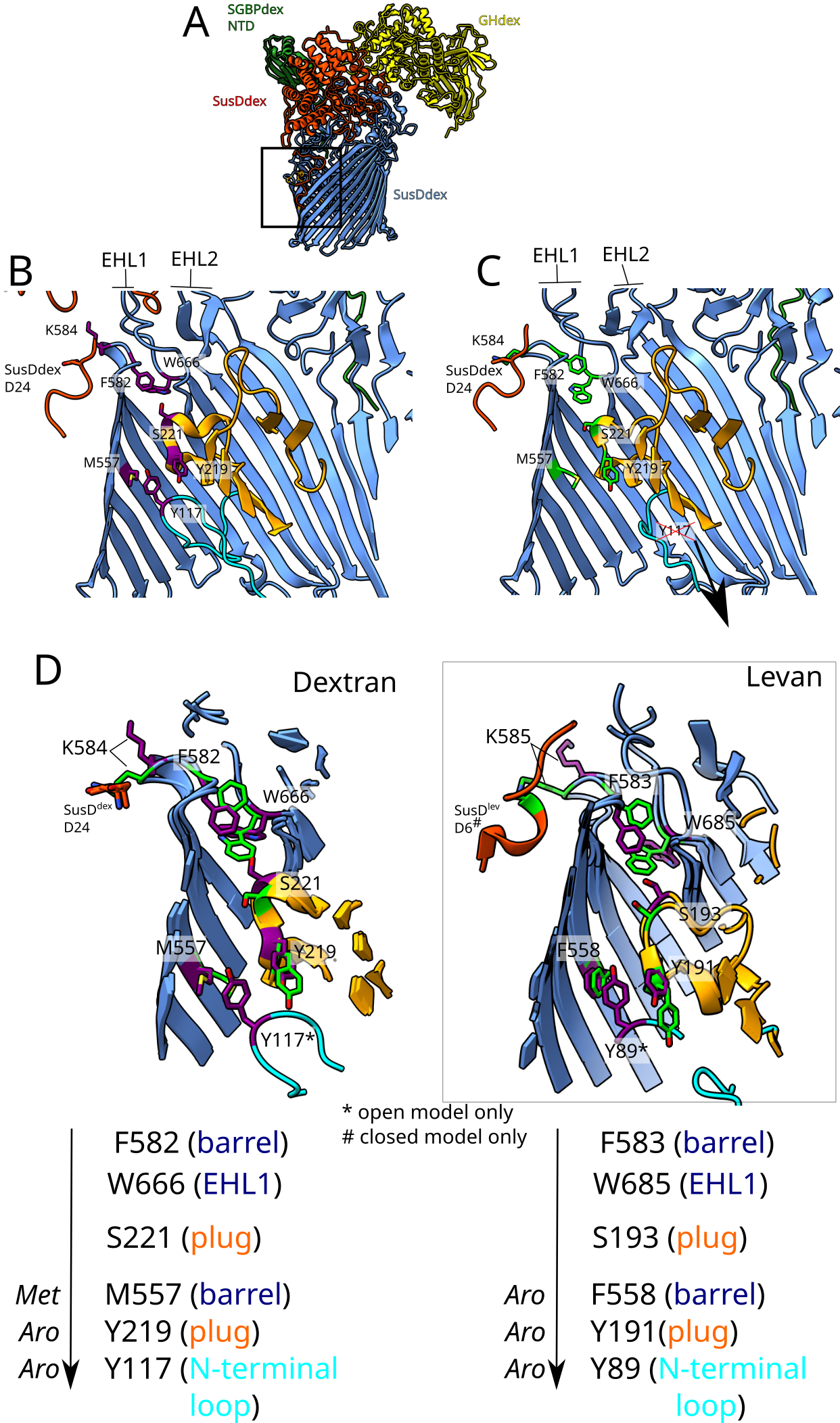

**Figure S16. The “aromatic lock” that facilitates release of the N-terminal TonB box of SusC^dex^.** The colour scheme is consistent with previous figures (SusC^dex^ blue, SusD^dex^ orange-red, SGBP^dex^ green and GH^dex^ yellow). The plug domain of SusC^dex^ is in orange while the N-terminal TonB box-containing loop is in cyan. **A.** The relevant area of the utilisome is highlighted. **B-C.** The open (**B**) and closed (**C**) models are shown in detail, with residues that undergo shifts between the models displayed. These include a methionine and two aromatics at the inner face of the barrel (Met557, Phe582, Trp666), the plug domain (Tyr219, Ser221), and the TonB box containing N-terminal loop (Tyr117, resolved only in open SusC^dex^). These residues are coloured purple (open state) or in lime green (closed state). Tyr117, which interacts with Met557 and Tyr219 in the open state, forms the aromatic lock, sequestering the TonB box inside the SusC^dex^ lumen. After closure of SusD^dex^, these interactions are broken, releasing the N-terminal loop into the periplasm for TonB to access the TonB box. **D.** Overlay of the open and closed states in the dextran and levan systems. Levan models include the active transporter, where no SusD^lev^ was modelled, but they are in the open state (PDB 8A9Y) and the inactive, closed SusD^lev^ model (PDB 8AA0). In the bottom panel, an arrow shows the direction of the allosteric signal from the top of SusC^dex^ (Phe582) to the N-terminal loop (Tyr117). The location of each shifted residue is in brackets.

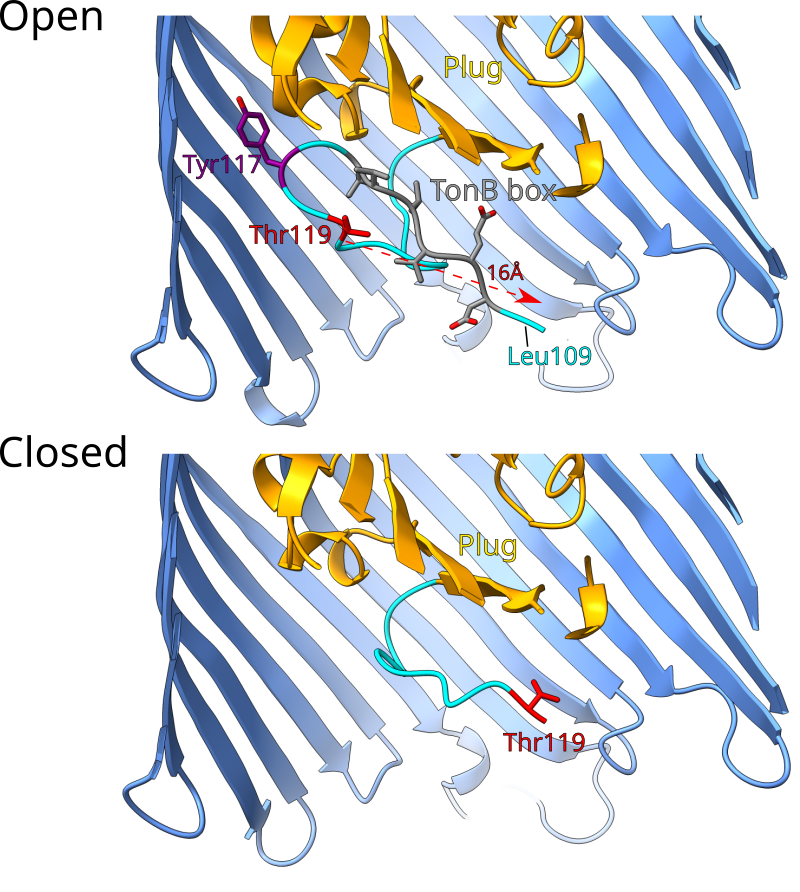

**Figure S17. SusC^dex^ N-terminus disordering after SusD^dex^ lid closure.** The N-termini of open and closed SusC^dex^ are compared, where the plug domain (residues 129-232) is orange and the N-terminal loop (109-128) is cyan. The TonB box residues (^110^DEVVVV^115^), predicted by homology with other SusCs, are in grey. Tyr117, involved in the aromatic lock (Figure S16), is in purple. The first residue resolved in both open and closed SusC^dex^ (Thr119) is coloured red, with a dashed arrow showing the difference in position (16 Å) between the two models. In the open model, the first resolved residue is also labelled (Leu109).

**Table S1. Data collection and refinement statistics for crystal structures of soluble constructs of the dextran utilisome surface lipoproteins.** Values in parentheses describe the last (highest resolution) shell.

| **Protein** | **GH^dex^** | | | **SGBP^dex^** | **SusD^dex^** | |
| --- | --- | --- | --- | --- | --- | --- |
| **Mutant, ligand** | **WT, apo** | **E360A, apo** | **E360A, +dex** | **Δ1-147, apo** | **WT, apo** | **WT, +dex** |
| **PDB** | 9SJG | 9SJH | 9SJF | 9SJI | 9SJK | 9SJJ |
|  | **Data collection** | | | | | |
| **Wavelength (Å)** | 0.898 | 0.979 | 0.898 | 0.898 | 0.898 | 0.98 |
| **Space group** | P 41 21 2 | P 41 21 2 | P 41 21 2 | C 1 2 1 | P 42 21 2 | I 2 2 2 |
| **Unit cell lengths a, b, c (Å)** | 93.70, 93.70, 331.61 | 92.86, 92.86, 330.68 | 92.22, 92.22, 328.03 | 148.32, 37.55, 65.62 | 119.50, 119.50, 176.98 | 87.00, 106.15, 138.34 |
| **Unit cell angles α, β, γ (°)** | 90, 90, 90 | 90, 90, 90 | 90, 90, 90 | 90, 100.35, 90 | 90, 90, 90 | 90, 90, 90 |
| **Resolution range** | 55.27 - 1.83  (1.83-1.80) | 330.68 - 2.10  (2.14-2.10) | 47.03 - 2.20  (2.25-2.20) | 36.48 - 1.70  (1.73-1.70) | 53.44 - 1.65  (1.68-1.65) | 67.29 - 1.70  (1.73-1.70) |
| **Total reflections** | 3455703  (183119) | 2217718  (110301) | 1816516  (60863) | 270053  (14440) | 7815948  (383132) | 1786941  (101033) |
| **Unique reflections** | 137723  (6704) | 85740  (4460) | 73184  (4453) | 39600  (2069) | 153600  (7518) | 70548  (3690) |
| **Multiplicity** | 25.1  (27.3) | 25.90  (24.70) | 24.80  (13.70) | 6.80  (7.00) | 50.9  (51.0) | 25.3  (27.4) |
| **Completeness (%)** | 100.00  (100.00) | 100.0  (100.0) | 100.00  (100.00) | 100.00  (100.00) | 100.00  (100.00) | 100.00  (100.00) |
| **mean(I) / σ(I)** | 13.40  (1.6) | 5.90  (1.20) | 7.70  (1.10) | 8.00  (0.70) | 19.3  (1.0) | 19.4  (1.3) |
| **Wilson B-factor** | 22.38 | 31.57 | 30.24 | 30.63 | 23.44 | 28.02 |
| **R_merge_** | 0.16  (2.27) | 0.90  (17.35) | 0.33  (2.32) | 0.09  (2.23) | 0.15  (4.26) | 0.10  (3.09) |
| **R_meas_** | 0.17  (2.31) | 0.92  (17.72) | 0.33  (2.40) | 0.10  (2.40) | 0.15  (4.31) | 0.10  (3.14) |
| **R_pim_** | 0.03  (0.44) | 0.18  (3.58) | 0.07  (0.63) | 0.04  (0.90) | 0.02  (0.60) | 0.02  (0.60) |
| **CC_half_** | 100.0  (81.0) | 99.0  (70.0) | 100.0  (51.0) | 100.0  (71.0) | 100.0  (83.0) | 100.0  (59.0) |
|  | **Refinement** | | | | | |
| **Reflections in refinement** | 137436 | 85542 | 73027 | 39575 | 153513 | 70548 |
| **Reflections in free set** | 6909 | 4260 | 3603 | 2011 | 7727 | 3544 |
| **R_work_** | 0.198 | 0.190 | 0.202 | 0.200 | 0.180 | 0.165 |
| **R_free_** | 0.229 | 0.230 | 0.262 | 0.270 | 0.206 | 0.191 |
| **FSC average** | 0.9695 | 0.9653 | 0.9623 | 0.9341 | 0.9763 | 0.9784 |
| **RMSD bonds** | 0.0094 | 0.0067 | 0.0083 | 0.0126 | 0.012 | 0.0135 |
| **RMSD angles** | 1.8150 | 1.791 | 1.885 | 2.214 | 2.077 | 2.075 |
| **Ramachandran favoured (%)** | 97.2 | 96.5 | 97.3 | 97.4 | 98.1 | 97 |
| **Ramachandran allowed (%)** | 2.8 | 3.5 | 2.7 | 2.6 | 1.8 | 2.6 |
| **Ramachandran outliers (%)** | 0 | 0 | 0 | 0 | 0.1 | 0.4 |
| **Rotamer outliers (%)** | 1.1 | 2.9 | 2.7 | 5.1 | 1 | 1.3 |
| **Clash score** | 10.4 | 12.9 | 7 | 8 | 1.6 | 3.5 |
| **MolProbity score** | 1.73 | 2.21 | 1.86 | 2.09 | 0.92 | 1.41 |

**Table S2. Characterisation of structure-based binding site variants of GH^dex^, SGBP^dex^ and SusD^dex^ via ITC.** The fold change (to the nearest integer) decrease in K_D_ was compared to the “wild-type” (WT) construct, described in brackets in the protein name, where each construct’s protein concentration was fixed at its calculated concentration (see Materials and methods). Ligand concentration was also fixed to the calculated value, so the wild-type values are slightly different from Table S4. Standard deviations (±) are shown in parentheses. Number of replicates refers to how many independent titrations were analysed. Titrations with no quantifiable binding are marked with NB. Example titrations are shown in Spreadsheet S2.

| **Protein** | **Dextran substrate (kDa)** | **Calculated dextran concentration (µM)** | **WT number of replicates** | **WT *K*_D_**  **(µM)** | **Variant number of replicates** | **Variant *K*_D_**  **(µM)** | **Decrease in *K*_D_ relative to WT (fold change)** |
| --- | --- | --- | --- | --- | --- | --- | --- |
| **(GH^dex^ D297A, E360A) W260A** | 1.5 | 1600 | 3 | 1.2  (±0.2) | 2 | 76.5  (±17.3) | 62 |
| **(SGBP^dex^)**  **W394A** | 500 | 12.5 | 3 | 2.8  (±0.4) | 3 | 13.2  (±4.4) | 5 |
| **(SGBP^dex^)**  **W433A** | 500 | 12.5 |  |  | 3 | 15.1  (±1.2) | 5 |
| **(SusD^dex^)**  **Y103A** | 1.5 | 1600 | 3 | 13.7 (±2.5) | 3 | 127.2  (±37.9) | 9 |
| **(SusD^dex^)**  **W284A** | 1.5 | 1600 |  |  | 3 | NB | NB |
| **(SusD^dex^)**  **F296A** | 1.5 | 1600 |  |  | 2 | NB | NB |

**Table S3. Data collection, map and real-space refinement statistics from cryo-EM of the dextran utilisome in the closed-closed (CC) and open-closed (OC) states**. Map resolution was estimated by the non-uniform refinement job in CryoSPARC (after FSC-mask auto-tightening and using a 0.143 cut-off) and supplied to Phenix for real-space refinement. Local resolution was estimated using the CryoSPARC job which implements a local windowed FSC method [7]. Models were refined using the final, octameric maps (see Figure S8C).

| **Data collection** | | | | |
| --- | --- | --- | --- | --- |
| **Category** | **Metric** | **OC**  **EMD-54945** | **CC**  **EMD-55024** | |
|  | **Number of movies** | 12,408 | | |
|  | **Magnification** | 105,000 | | |
|  | **Defocus range (µm)** | -1.0 to -2.0 | | |
|  | **Microscope** | TFS Krios | | |
|  | **Voltage (kV)** | 300 | | |
|  | **Electron dose (e^-^/Å^2^)** | 23.1 | | |
|  | **Image detector** | Gatan K3 | | |
| **Processing** | | | | |
| **Number of particles** | **Picked** | 4,556,980 | | |
|  | **After 2D classification** | 481,934 | | |
|  | **Final 3D refinement** | 23,484 | 116,104 | |
|  | **Recommended contour level** | 0.025 | 0.015 | |
|  | **Symmetry imposed** | C1 | C1 | |
|  | **Pixel spacing (Å/pixel)** | 0.831 | | |
|  | **Map resolution (Å)**  **(FSC threshold 0.143)** | 2.9 | | 2.49 |
| **Local resolution estimation (Å)**  **(FSC threshold 0.5)** | **Range** | 1.78 to 48.49 | 2.19 to 39.53 | |
|  | **25^th^ percentile** | 3.76 | 2.81 | |
|  | **Median** | 6.92 | 4.35 | |
|  | **75^th^ percentile** | 8.35 | 6.14 | |
| **Refinement** | | | | |
| **Category** | **Metric** | **OC** | **CC** | |
| **PDB** |  | **9SJE** | **9SM2** | |
| **Composition (#)** | **Chains** | 11 | 11 | |
|  | **Atoms** | 32922 (Hydrogens: 0) | 32876 (Hydrogens: 0) | |
|  | **Residues** | Protein: 4131 Nucleotide: 0 | Protein: 4124 Nucleotide: 0 | |
|  | **Water** | 0 | 0 | |
|  | **Ligands** | GLC: 35 | GLC: 36 | |
| **Bonds (RMSD)** | **Length (Å) (# > 4σ)** | 0.004 (0) | 0.003 (2) | |
|  | **Angles (°) (# > 4σ)** | 0.816 (12) | 0.691 (13) | |
| **MolProbity** | **MolProbity score** | 1.56 | 1.37 | |
|  | **Clash score** | 1.99 | 1.79 | |
| **Ramachandran plot (%)** | **Outliers** | 0.70 | 0.58 | |
|  | **Allowed** | 4.50 | 4.02 | |
|  | **Favored** | 94.80 | 95.40 | |
| **Rama-Z** | **Whole (N = 4115 / 4108)** | -0.19 (0.13) | 0.14 (0.13) | |
|  | **Helix** | -1.21 (0.15) | -0.62 (0.16) | |
|  | **Sheet** | 1.04 (0.15) | 1.21 (0.15) | |
|  | **Loop** | -0.23 (0.14) | -0.13 (0.14) | |
| **Side Chain Quality** | **Rotamer outliers (%)** | 2.07 | 1.41 | |
|  | **Cß outliers (%)** | 0.00 | 0.00 | |
| **Peptide Plane (%)** | **Cis proline / general** | 1.4 / 0.0 | 1.4 / 0.0 | |
|  | **Twisted proline / general** | 0.0 / 0.0 | 0.0 / 0.0 | |
| **CaBLAM outliers (%)** |  | 2.83 | 2.17 | |
| **ADP (B-factors)** | **Iso/Aniso (#)** | 32922 / 0 | 32876 / 0 | |
|  | **Protein (min/max/mean)** | 14.39 / 236.92 / 106.65 | 28.15 / 248.56 / 102.54 | |
|  | **Ligand (min/max/mean)** | 20.00 / 166.78 / 98.05 | 20.00 / 147.15 / 90.05 | |
| **Occupancy** | **Mean** | 1.00 | 1.00 | |
|  | **occ = 1 (%)** | 100.00 | 100.00 | |
|  | **0 < occ < 1 (%)** | 0.00 | 0.00 | |
|  | **occ > 1 (%)** | 0.00 | 0.00 | |
| **Box Dimensions (Å)** | **Lengths** | 152.90, 174.51, 177.00 | 154.57, 178.66, 181.16 | |
|  | **Angles** | 90.00, 90.00, 90.00 | 90.00, 90.00, 90.00 | |
| **Resolution (Å)** | **Supplied** | 2.9 | 2.5 | |
|  | **d99 (full/half1/half2)** | 3.5 / --- / --- | 3.1 / --- / --- | |
|  | **d model** | 3.3 | 3.0 | |
|  | **d FSC model (0/0.143/0.5)** | 2.8 / 2.9 / 3.2 | 2.5 / 2.6 / 3.1 | |
| **Map** | **min / max / mean** | -0.09 / 0.19 / 0.00 | -0.07 / 0.19 / 0.00 | |
| **Model vs. Data** | **CC (mask)** | 0.88 | 0.87 | |
| **(Cross-correlation)** | **CC (box)** | 0.76 | 0.77 | |
|  | **CC (peaks)** | 0.66 | 0.66 | |
|  | **CC (volume)** | 0.87 | 0.87 | |
|  | **Mean CC for ligands** | 0.72 | 0.81 | |

**Table S4. Binding affinity (*K*_D_) for purified recombinant dextran utilisome surface lipoproteins when titrated with dextrans ranging from 1.5 to 500 kDa.** This analysis was performed by fixing the protein concentration to the calculated amount and N (stoichiometry) to 1, while allowing the software to change the dextran concentration from the calculated value to fit the data (the expected values given to the software are listed in column 3). To represent the variation in each set of replicates, standard deviations (±) are shown. Example titrations are shown in Spreadsheet S2.

| **Protein** | **Dextran substrate (kDa)** | **Calculated dextran concentration (µM)** | **Number of replicates** | ***K*_D_**  **(µM)** |
| --- | --- | --- | --- | --- |
| **GH^dex^ E360A** | **1.5** | 1600 | 3 | 2.54 (±0.22) |
|  | **3.5** | 800 | 3 | 1.29 (±0.13) |
|  | **10** | 250 | 3 | 2.81 (±0.04) |
|  | **20** | 100 | 3 | 2.11 (±0.26) |
|  | **40** | 50 | 5 | 3.51 (±1.16) |
|  | **150** | 25 | 3 | 2.84 (±0.73) |
|  | **500** | 12.5 | 3 | 2.33 (±1.32) |
| **GH^dex^ D297A, E360A** | **1.5** | 1600 | 3 | 2.01 (±0.58) |
|  | **3.5** | 800 | 3 | 3.01 (±0.33) |
|  | **500** | 12.5 | 3 | 3.72 (±1.58) |
| **SGBP^dex^** | **1.5** | 1600 | 5 | 6603.24 (±9039.12) |
|  | **3.5** | 800 | 3 | 25.97 (±2.25) |
|  | **10** | 400 | 3 | 16.23 (±0.97) |
|  | **20** | 400 | 3 | 8.43 (±0.20) |
|  | **40** | 400 | 3 | 6.55 (±0.86) |
|  | **150** | 25, 30 | 3 | 6.10 (±1.22) |
|  | **500** | 12.5 | 3 | 2.76 (±0.45) |
| **SGBP^dex^ Δ1-147** | **1.5** | 1600 | 3 | 101.90 (±14.9) |
|  | **3.5** | 800 | 3 | 32.83 (±3.73) |
|  | **10** | 800 | 3 | 20.07 (±2.66) |
|  | **20** | 200 | 3 | 11.57 (±0.65) |
|  | **40** | 100 | 3 | 6.71 (±0.76) |
|  | **150** | 50 | 3 | 3.66 (±0.58) |
|  | **500** | 10, 12.5 | 3 | 3.23 (±2.49) |
| **SusD^dex^** | **1.5** | 1600 | 3 | 15.57 (±1.92) |
|  | **3.5** | 800 | 3 | 12.56 (±4.01) |
|  | **10** | 250 | 3 | 13.77 (±0.06) |
|  | **20** | 100 | 3 | 7.69 (±1.33) |
|  | **40** | 50 | 3 | 14.70 (±2.63) |
|  | **150** | 25 | 3 | 15.53 (±2.57) |
|  | **500** | 12.5 | 3 | 18.70 (±0.70) |

**Table S5. Stoichiometry analysis of dextran binding proteins.** Using a one set of sites binding model, the N value (number of binding sites) was allowed to vary to fit the data while keeping the supplied dextran concentration fixed. This is different from the previous analysis (Table S4), where N was fixed to equal 1 and the dextran concentration was varied. Three independent runs were analysed using this method and the N values were then averaged. Values marked with asterisks were deemed untrustworthy due to the software’s modelling function hitting a lower bound for ΔH and these values were not averaged. See also the Supplementary Discussion.

| **Dextran binding protein** | **Dextran 1.5 (N)** | **Mean** | **Dextran 3.5 (N)** | **Mean** |
| --- | --- | --- | --- | --- |
| **GH^dex^ E360A** | 1.98, 2.25, 2.13 | 2.12 | 0.920, 0.934, 0.976 | 0.94 |
| **SGBP^dex^** | 0.630*, 0.606, 0.336*, 0.483* | N/A | 1.26, 1.26, 1.17 | 1.23 |
| **SGBP^dex^ Δ1-147** | 1.83, 1.36, 1.34 | 1.51 | 0.556*, 0.758, 1.36 | N/A |
| **SusD^dex^** | 1.01, 1.03, 1.00 | 1.01 | 0.492, 0.507, 0.463 | 0.49 |

**Table S6. Descriptions of Movies S1 and S2**

| **Movie** | **Description** |
| --- | --- |
| Movie S1. The closing of SusD^dex^ and SGBP^dex^ supplying  glycans to GH^dex^ | An accompaniment to Figure S8. A 20-frame movie was generated by CRYOSPARC 3D variability on a consensus stack of ~270k particles. This movie shows the SusD^dex^ in motion from its open to closed state. The C-terminal domain (CTD) of SGBP^dex^, which is poorly resolved in 3D reconstructions used for model building, is observed at the beginning and the end of this motion. At the beginning, the CTD contacts the GH^dex^ binding site and a bridge of density is observed (44-45s); this may reflect how glycans are passed from SGBP^dex^ to GH^dex^. As the SusD^dex^ closes, SGBP^dex^ simultaneously shifts away from GH^dex^ and appears to be more resolved resting upon the top of closed SusD^dex^ (43-44s). |
| Movie S2. Dextran utilisome binding sites in the closed-closed model. | An accompaniment to Figure 5. Movie generated in ChimeraX. Binding sites for GH^dex^, SusD^dex^ and SusC^dex^ are shown in sequence, with bound IMO6, IMO8 and IMO4 (dextran ligands), respectively. Residues within 5Å of each dextran ligand are displayed as grey sticks, while IMO are displayed in magenta. SusC^dex^ is cornflower blue, while its plug domain (residues 129-232) is coloured orange. SusD^dex^ is orange-red, GH^dex^ is yellow and SGBP^dex^ is forest green. Only the N-terminal domain of SGBP^dex^ (non-carbohydrate-binding, Ig-like) was resolved sufficiently for modelling. |

**Supplementary discussion**

**GH^dex^ crystal structure.**

If catalysis is still occurring in GH^dex^ E360A, it is puzzling why ligand was not resolved at subsite -1 (see Figure 2). Perhaps the cleavage site is more promiscuous and can occur between subsites -1 and -2. It is worth noting that the three resolved dextran residues are almost identical to a crystal structure of a triose bound to the *Flavobacterium johnsonia* GH66 dextranase, which is the most similar structural homologue in the PDB (with ~48% sequence identity to GH^dex^) [8]. The authors of the *Tp*^dex^ crystal structure noted that between the two molecules of the asymmetric unit, W376 either occluded subsite +1 or provided a “side wall” of the catalytic cleft , and this may be responsible for the expulsion of hydrolysed glycans (Figure S2, panel C) [2]. However, this was not observed in either crystal and cryo-EM structures of GH^dex^, with Trp362 in the side wall conformation in both apo and liganded states.

**Estimation of number of SLP binding sites.**

To infer the number of binding sites of these dextran-binding proteins, the two of the smallest dextrans available from Pharmacosmos (technical grade dextran 1.5 and 3.5) were compared in a separate ITC data analysis where the N value was allowed to vary to estimate dextran-to-protein stoichiometry (Table S5). We hypothesised that dextran 1.5 was small enough (DP = 6-10) that it could not bind more than one binding site simultaneously. These results were then compared to dextran 3.5 (DP = 16-22), and N values were compared. This suggested SusD^dex^ had one binding site for dextran 1.5 (derived N = 1.01) and half this for dextran 3.5 (N = 0.49), indicating that glycans of DP 6-10 are typical substrates for SusD^dex^. This agreed with predictions that SusD^dex^ would contain one binding site based on homology and previous structural data [9–11].

For GH^dex^ E360A, titrations of dextran 1.5 (N = 2.12) indicated that two dextran molecules were bound per protein. Conversely, dextran 3.5 gave an average value of 0.94 indicating that this dextran (DP ~20) may be large enough to span two putative binding sites in GH^dex^. In the cryo-EM structure of the substrate-loaded levan utilisome, as well as the crystal structure of GH^lev^, two sites were observed on GH^lev^: one forming the active site and another on top of the catalytic domain [11]. These sites did not bind the same levan ligand simultaneously, but this may be possible with longer ligands in solution. However, unlike GH^lev^, the crystal or cryo-EM structures of GH^dex^ did not suggest two distinct binding sites. SusG from the starch PUL also contains two binding sites, but this is provided by a distinct CBM domain [12]. These GHs belong to distinct families to GH^dex^; GH13 and GH32 respectively. The second closest structural homologue in the PDB, GH66 family member *Tp*^dex^, had only one binding site [2].

SGBP^dex^ is homologous to SusF of the starch PUL, which was identified to have three binding sites for glycans [3]. By contrast, the structure of SGBP^lev^ is divergent with only one binding site [11]. N value analysis was unreliable for SGBP^dex^, likely due to the presence of several distinct binding sites and its low affinity for small dextrans (Table S5).

**SGBP^dex^ Δ1-147 crystal structure.**

In the crystal structure of SGBP^dex^ Δ1-147, a loop at the top surface of CBM2 contains a cis peptide bond between Ala306 and Asp307 (Figure S4), which is rare for bonds that do not contain prolines (cis non-Pro) [13]. This bond is well supported by electron density (Figure S4). Cis non-Pro motifs have been identified to be important for biological activity, including ligand binding, dimerisation and catalysis [14–16]. Genuine cis non-Pro bonds are also relatively more common in carbohydrate-active proteins, in which they are overrepresented compared to non-carbohydrate binding proteins [16]. In SGBP^dex^, conformational restriction by an upstream proline (Pro303) may influence this bonding. There are reports of downstream prolines reducing the barrier to cis/trans rotation in the peptide preceding the proline [16,17], but it is unclear if upstream prolines also have an effect. Ala307/Asp308 is surrounded by candidate surface-exposed dextran-binding residues including an angled surface of Trp281-Phe318 and Trp304. Somewhat similarly, in SusF, CBM2 residues Trp287 and Trp330 form an aromatic platform that binds starch, behind which Thr289-Pro290 form a cis-Pro peptide bond [3]. The SGBP^dex^ cis peptide between Ala306-Asp307 may support dextran binding by projecting Asp307 towards a glycan cooperatively bound by the proximal Trp281-Phe318 aromatic surface residues, which are spatially analogous to confirmed binding residues Trp287 and Trp330 in SusF CBM2. Asp307 may stabilise bound dextran by hydrogen bonding. Additionally, dextran binding to CBM2 could instigate cis-to-trans isomerisation, driving the movement of SGBP^dex^ towards GH^dex^ seen in cryo-EM (Figure S8). There are also other examples of functional isomerisation of cis non-Pro peptide bonds. Concanavalin A, a lectin, is known to be in equilibrium between locked and unlocked conformational states. Based on its crystal structure, this was proposed to be caused by the binding of a metal ion, which drives a trans-to-cis isomerisation of an Ala-Asp peptide bond [18]. Binding of the glycan methyl-α-D-mannose alone was also observed to lock metal-free concanavalin A via peptide bond isomerisation [18].
